# Antibiotic heteroresistance generated by multi-copy plasmids

**DOI:** 10.1101/2022.08.24.505173

**Authors:** JCR Hernandez-Beltran, J Rodríguez-Beltrán, B Aguilar-Luviano, J Velez-Santiago, O Mondragón-Palomino, RC MacLean, A Fuentes-Hernández, A San Millán, R Peña-Miller

## Abstract

Heteroresistance – in which a clonal bacterial population contains a cell subpopulation with higher resistance to antibiotics than the main population – is a growing clinical problem that complicates susceptibility determination and threatens therapeutic success. Despite the high prevalence of heteroresistance in clinical settings, the underlying genetic mechanisms that stably maintain heterogeneous bacterial populations are poorly understood. Using fluorescence microscopy, single-cell microfluidics, and quantitative image analysis, we show that random replication and segregation of multicopy plasmids produce populations of bacterium *Escherichia coli* MG1655 in which cells with low-and high-plasmid copy numbers stably co-exist. By combining stochastic simulations of a computational model with high-throughput single-cell measurements of *bla*_TEM-1_ expression, we show that copy number variability confers the bacterial population with transient resistance to a lethal concentration of a *β* -lactam antibiotic. Moreover, this surviving, high plasmid copy minority is capable of regenerating a heterogeneous bacterial population with low and high plasmid copy numbers through segregational instability, rapidly alleviating the fitness burden of carrying large numbers of plasmids. Our results provide further support for the tenet that plasmids are more than simple vehicles for horizontal transmission of genetic information between cells, as they can also drive bacterial adaptation in dynamic environments by providing a platform for rapid amplification and attenuation of gene copy number that can accelerate the rate of resistance adaptation and can lead to treatment failure.

## Introduction

The evolution and spread of antimicrobial resistance in clinical pathogens represent a major public health problem that threatens to become a global crisis.^1^ In general, drug resistance is considered to be the consequence of stable genetic mutations or the acquisition of antibiotic resistance genes through horizontal gene transfer.^2^ However, treatment failure can also result from the presence of subpopulations of bacterial cells with higher levels of resistance than those of the rest of the population.^3^ This phenomenon is known as heteroresistance^4, 5^ and has been identified in diverse bacterial species and in a wide range of antimicrobial classes.^6–8^

Previous studies have shown that increased tolerance to antimicrobial substances can be achieved through a subset of dormant cells, known as persisters, that survive drug exposure and resume growth once the antibiotic is withdrawn.^9^ Moreover, there are several genetic and metabolic mechanisms that generate subpopulations with differing degrees of drug tolerance,^10, 11^ for instance through the heterogeneous production of drug-degrading enzymes^8, 12^ or signaling molecules.^13^ Heterogeneous drug susceptibility within a population can also arise from the stochastic expression of genes encoding intrinsic antibiotic-resistance mechanisms, notably efflux pumps.^14, 15^

Rapid adaptation to antibiotics can also be achieved through genomic duplications that increase the dosage of known drug-resistance genes,^8, 16, 17^ for instance through amplification of efflux pump operons^18, 19^ or genes encoding drug-modifying enzymes.^20, 21^ Laboratory studies have shown that genomic amplifications scale up with the strength of the selective pressure,^16^ and are unstable in the absence of selection due to the fitness burden associated with the duplication of large chromosome regions.^16, 22, 23^

In the clinic, heteroresistance due to spontaneous tandem gene amplification has been proposed as a plausible cause of treatment failure,^24^ with the incidences likely to be underestimated due to the intrinsic limitations of standard microbiology assays.^25^ A recent large-scale analysis of heteroresistant clinical isolates found a high incidence of genomic amplifications that increased resistance to multiple antibiotics.^26^ Interestingly, whole-genome sequencing revealed that, while some duplications occurred in large chromosomal regions containing known drug resistance genes, a considerable fraction of sequence amplifications were found in plasmids.

Plasmids are DNA molecules that replicate independently of the chromosome and play an essential role in the dissemination of resistance genes among clinically important pathogens.^27^ Crucially, plasmids can be present in multiple copies per cell, from a few copies to dozens for high-copy plasmids. Although some plasmids can be transferred horizontally, thus spreading resistance genes between bacterial hosts, a large fraction of plasmids are non-conjugative and are carried in multiple copies per cell.^28^ A recent clinical study showed that a large fraction of pathogenic *Escherichia coli* isolates carry small ColE1 plasmids.^29^ The number of plasmids carried by each cell is a key driver of virulence^30^ and horizontal gene transfer.^31^ Furthermore, cells within a biofilm contain high plasmid copy numbers and therefore have elevated transcription of antibiotic resistance genes.^32^

For multicopy plasmids lacking active partitioning or postsegregational killing mechanisms,^33^ segregation occurs randomly upon division, with the probability of a plasmid being inherited to a given cell following a binomial distribution.^34–36^ The interaction between replication and segregation, and the complex population dynamics this produces^37, 38^ is known to enhance bacterial adaptation to novel environmental conditions,^39^ as well as to determine the repertoire of genes carried in plasmids^40^ and their stability in the absence of selection.^41, 42^ Moreover, recent studies have shown that multicopy plasmids can accelerate bacterial adaptation,^43^ for instance by promoting intracellular genetic diversity^44^ and increasing the probability of the appearance of beneficial mutations and subsequently amplifying mutant gene expression.^45^

In addition to amplifying gene dosage, an increase in copy number is also associated with a decrease in the probability of plasmid loss and with a higher metabolic burden.^46^ A consequence of this tradeoff is that plasmid replication is subject to two conflicting levels of selection:^35, 47, 48^ plasmids that overreplicate have a higher chance of overcoming segregational loss and becoming fixed in descendant cells, but cells with more plasmid copies have a lower probability of becoming fixed in the population. As a result, plasmid control is a tightly regulated process^49^ that depends on the host’s genetic^50^ and physiological state,^51^ as well as on the extracellular environmental conditions.^52, 53^ For high-copy plasmids, however, replicative noise emerges as intracellular selection favors overreplication, thereby relieving intracellular selection for precise copy number control.^35^

We hypothesized that heteroresistance to a *β* -lactam antibiotic can emerge from cell-to-cell differences in plasmid copy number (PCN) in otherwise genetically identical cells. In the present study, we used a combination of single-cell and population-level experiments to show that encoding drug resistance genes in multicopy plasmids is beneficial in rapidly changing environments, as it enables bacterial communities to implement a reversible phenotypic tolerance mechanism based on the stable co-existence of susceptible and resistant cells. These experimental results were recapitulated by a computational model in which plasmid copy number variability was the main driver of cell-to-cell differences.

## Results

### Environmental modulation of PCN distributions in bacterial populations

To investigate the distribution of plasmids in bacterial populations, we used an experimental model system consisting of *E. coli* MG1655 carrying pBGT, a ColE1-like plasmid containing a GFP fluorescent marker (*eGFPmut2*) and *bla*_TEM-1_, a gene that encodes a TEM-1 *β* -lactamase, which inactivates *β* -lactam antibiotics by hydrolyzing the *β* -lactam ring.^54^ *β* -lactam resistance genes are generally located on plasmids and, in particular, TEM-1 has a plasmid origin, with more than two-hundred TEM *β* -lactamase variants descending from this allele recorded.^55^ We denote the strain carrying this well-characterized,^45, 56^ non-conjugative, and multicopy plasmid as MG/pBGT (average copy number=19.12, s.d.= 1.53; Figure 1A-C).^45^

**Figure 1.**
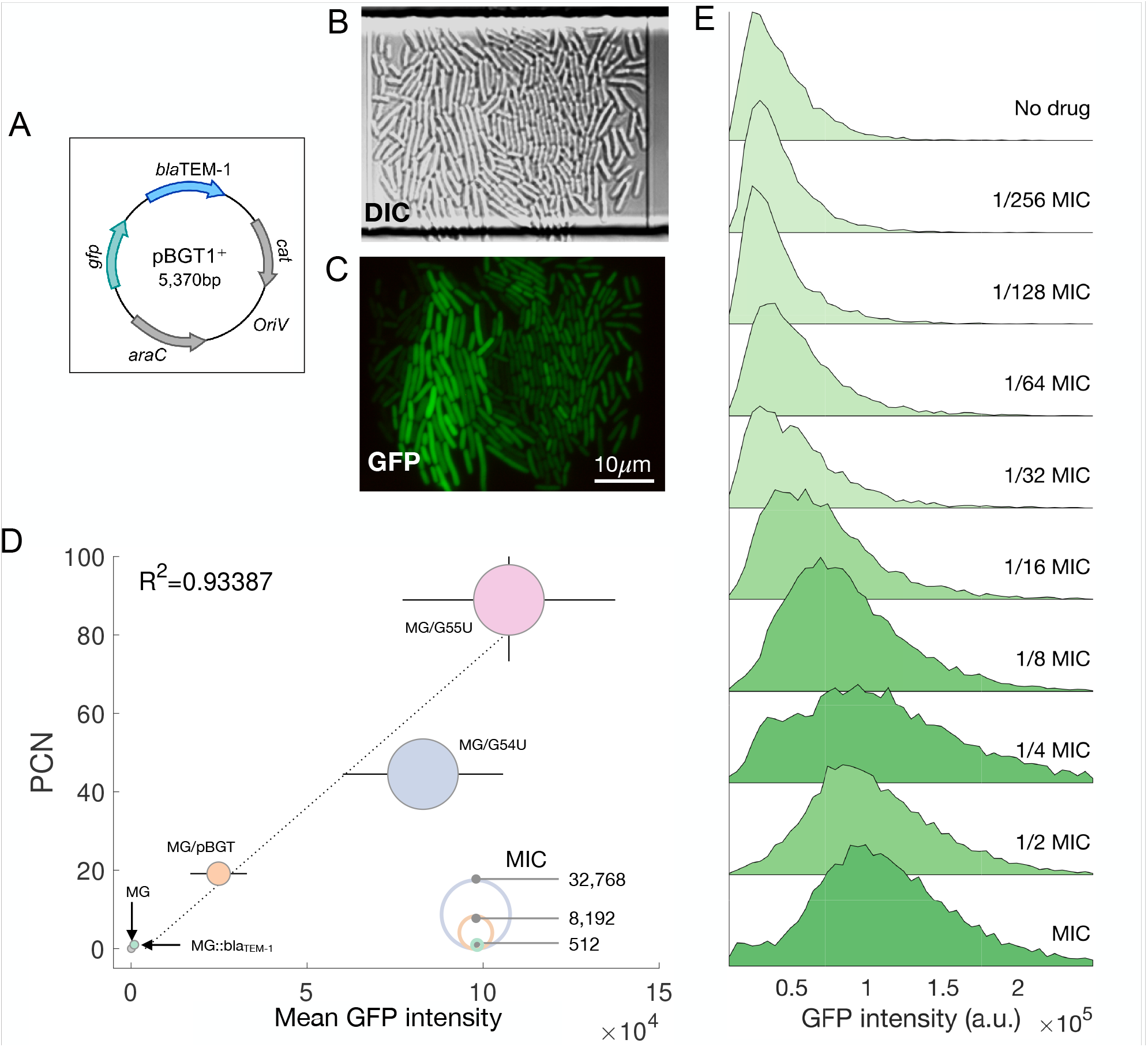
Experimental model system. A) Schematic representation of plasmid pBGT encoding *bla*_TEM-1_ (in blue) and *eGFPmut2* (in green). The reading frames for genes are represented with arrows, with arrowheads indicating the direction of transcription. B) DIC microscopy image of a plasmid-bearing population (MG/pBGT). C) Fluorescence microscopy image of this population shows high levels of GFP heterogeneity between cells. D) Mean fluorescence and mean plasmid copy number are positively correlated in bacterial populations. The circle’s diameter is proportional to each strain’s drug resistance level. E) Fluorescence distributions of MG/pBGT exposed to a range of ampicillin concentrations.

As a control, we used a fluorescently tagged strain carrying a chromosomally encoded *bla*_TEM-1_, which we term MG:GT. Moreover, to explore the association between PCN and fluorescence, we also used strains obtained in a previous experimental evolution study,^45^ with mutations in the origin of replication (Table S1) that result in a high mean PCN, with correspondingly high fluorescence intensity and elevated drug resistance compared with MG/pBGT.

In a recent study, direct, fluorescent-reporter-based measurement of PCN, promoter activity, and protein abundance at single-cell resolution revealed a positive correlation between PCN and protein expression.^57^ In our experimental system, we similarly observed a correlation between PCN measured by qPCR^39^ and fluorescence intensity quantified using a fluorescence spectrophotometer (*R*^2^ = 0.9387; Figure 1D). To validate the correlation between PCN and GFP in our system, we sorted the plasmid-bearing cell population according to GFP intensity into clusters with low, medium, and high fluorescence and confirmed the positive correlation between fluorescence and mean PCN estimated by qPCR (*R*^2^ = 0.879; Figure S1).

To measure the effect of the strength of antibiotic selection pressure on the distribution of PCN, we exposed a population of MG/pBGT cells to a range of ampicillin (AMP) concentrations, and used flow cytometry to measure GFP abundance in single cells. We found that the mean GFP abundance increased with the strength of selection (Figure 1E), and that the coefficient of variation for the PCN distribution decreased as a function of drug concentration (*R*^2^ = 0.593, p-value< 0.01; Figure S3). When the same experiment was repeated with MG:GT cells, mean fluorescence and its coefficient of variation remained constant accross all AMP concentrations (*R*^2^ = 0.052, p-value*>* 0.5; Supplementary Figure S2).

### The rapid increase in PCN is a population-level effect

The increase in fluorescence that we observed in response to AMP could be explained by a uniform increase in resistance levels in all cells in the population (e.g. by increasing the rate of plasmid replication), or by cell-to-cell heterogeneity in resistance levels (e.g. pre-existing copy number variability in the population). Because population-level experiments cannot be used to contrast both hypothesis, we measured the response of individual MG/pBGT cells to AMP exposure using a microfluidic chemostat and fluorescent microscopy (Methods). Using this set-up, we followed the life history of individual plasmid-bearing cells exposed to an antibiotic ramp of linearly increasing AMP concentration.

In these experiments, the longest surviving MG/pBGT cells were those with high fluorescence before antibiotic exposure (Figure 2A and Movie S1). Crucially, the fluorescence of individual cells remained constant throughout the antibiotic ramp (Figure 2B), suggesting that the population-level increase in mean GFP is a consequence of the antibiotic killing low PCN cells, and not the result of individual cells upregulating plasmid replication or *bla*_TEM-1_ expression (see also Figure S4).

**Figure 2.**
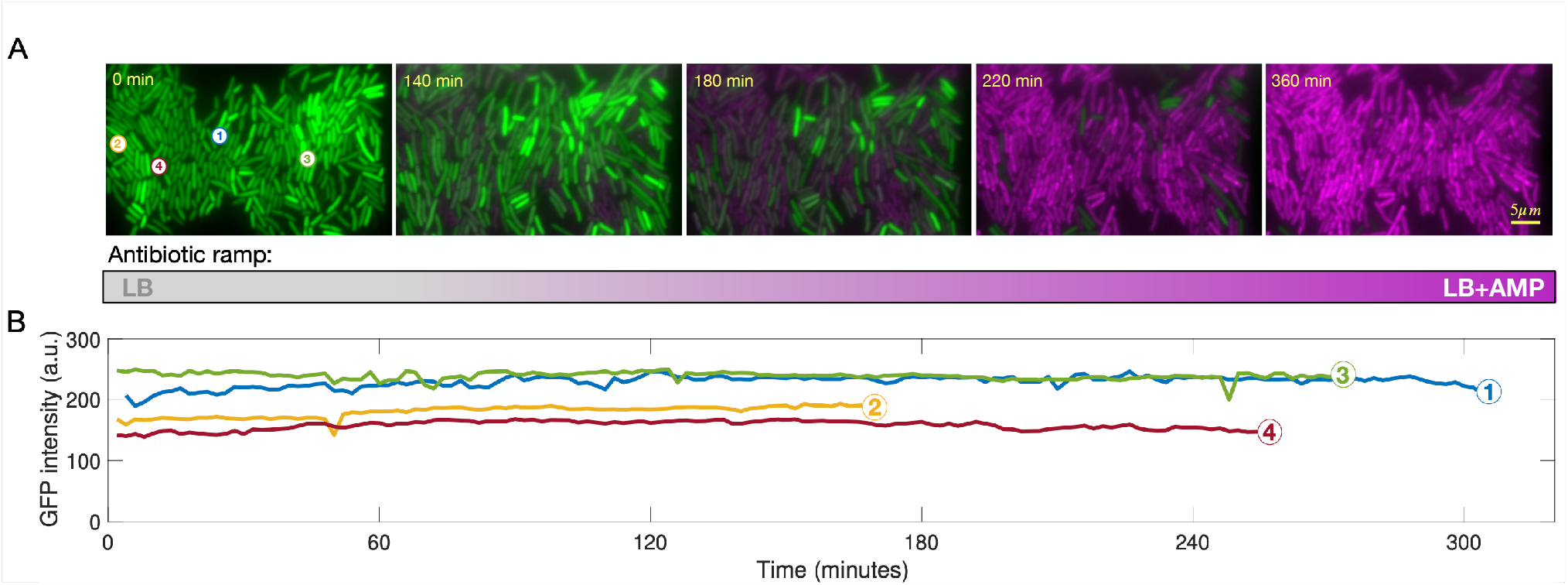
Microfluidics antibiotic ramp. A) Time-lapse movie showing a population of MG/pBGT cells exposed to increasing antibiotic concentrations (green channel: GFP fluorescence; magenta: red fluorescent dye, as a reporter of cell death). Note how cells with increased tolerance to the antibiotic have high levels of GFP fluorescence at the start of the antibiotic ramp, suggesting that fluorescence intensity and tolerance to the antibiotic are correlated at the single-cell level. B) Representative time-series of mean fluorescent intensity of individual cells suggest that TEM-1 expression remains constant throughout the experiment. Circles represent the moment each cell dies.

### PCN variability enhances the survival of bacterial populations exposed to fluctuating selection

To determine if between-cell differences in drug resistance produced heteroresistance at a population-level, we examined the response to AMP of 88 clonal populations of plasmid-bearing strains with different mean PCNs (strains MG/pBGT, MG/G54U and MG/G55U, with 19, 44, and 88 plasmid copies, respectively). When the cultures reached exponential growth, ∼ 1% of each population was transferred to an environment containing replenished media and a lethal AMP concentration (Figure 3A). After 30 minutes, a sample was transferred back to drug-free medium. This sampling process was repeated every 30 minutes and, for each duration of drug exposure, we counted the number of replicates showing growth after 24h.

**Figure 3.**
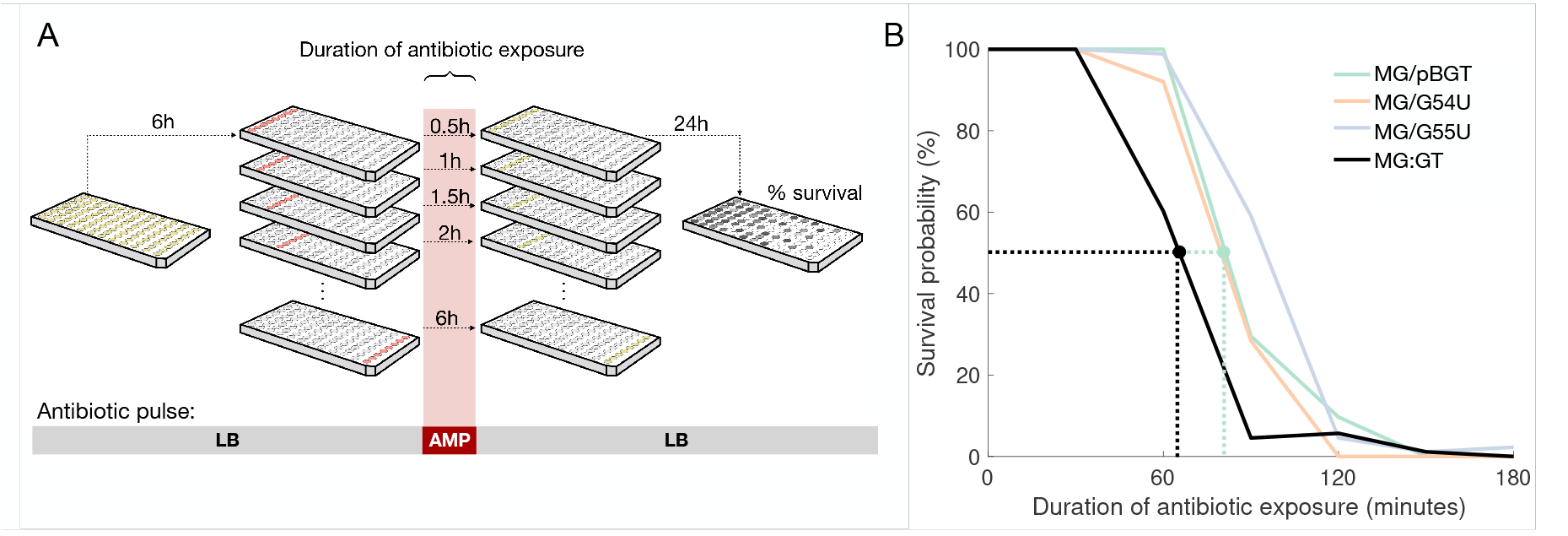
Multi-copy plasmids increase survival to antibiotic pulses. A) Diagram of a survival assay consisting in transiently exposing 88 populations of MG:GT and MG/pBGT to a lethal concentration of AMP. By sampling each population every 30 minutes and transferring it to drug-free media, we estimate the probability of survival of each strain based on the percentage of surviving populations after 24 hours of growth in drug-free media. B) Kaplan-Meier plot comparing survival probabilities as a function of the time exposed to a lethal ampicillin concentration (with MIC determined separately for each strain). Dotted lines represent the duration of drug exposure that results in a 50% survival probability (MG:GT in black, MG/pBGT in green).

Relative to MG:GT, all plasmid-bearing populations exhibited increased survival of fluctuating selection (Figure 3B; log-rank test, p-value*<* 0.005). For instance, the probability of survival after 90 minutes of AMP exposure was *>* 50% for all plasmid-bearing strains, whereas *<* 5% of the MG:GT replicate populations survived. It should be noted that the lethal drug concentration was determined independently for each strain (see Table S1 for MICs used). For each strain, we estimated the duration of drug exposure such that the probability of survival was 50% (60 min for MG:GT at 2 *mg/mL* AMP, and 80 min at 32 *mg/mL* AMP for MG/pBGT; dotted lines in Figure 3B). We refer to exposure to this concentration and duration as a *semi-lethal pulse*.

To confirm that the increased tolerance to a semi-lethal pulse presented by plasmid-bearing strains was not a consequence of a decrease in growth rate associated with the metabolic burden inherent to carrying plasmids (rather than selection of a subpopulation with more copies of *bla*_TEM-1_), we performed a survival assay for MG/pBGT in the presence of 256 *µg/L* of the *β* -lactamase inhibitor sulbactam. As expected, fluorescence remained constant independently of the ampicillin concentration, and only one out of eight populations survived exposure to 2 *mg/mL* of AMP (Figure S5).

### Quantifying heterogeneity in *bla*_TEM-1_ expression and survival after a semi-lethal pulse

In a microfluidic experiment, MG/pBGT and MG:GT populations were exposed separately to a semi-lethal pulse of AMP, with the critical dose and duration of the antibiotic pulse determined independently for each strain (Figure 4A). We acquired time-series of the fluorescent intensity of individual cells, recorded division events, and estimated the duplication rates of 5, 810 lineages for MG/pBGT and 1, 077 MG:GT lineages, respectively obtained from 46 and 8 separate microfluidic chambers (see Supplementary Movies 2 and 3 for sample time-lapse movies). The criteria for including a single-cell lineage in the analysis was that they were observed for a period spanning the antibiotic pulse.

**Figure 4.**
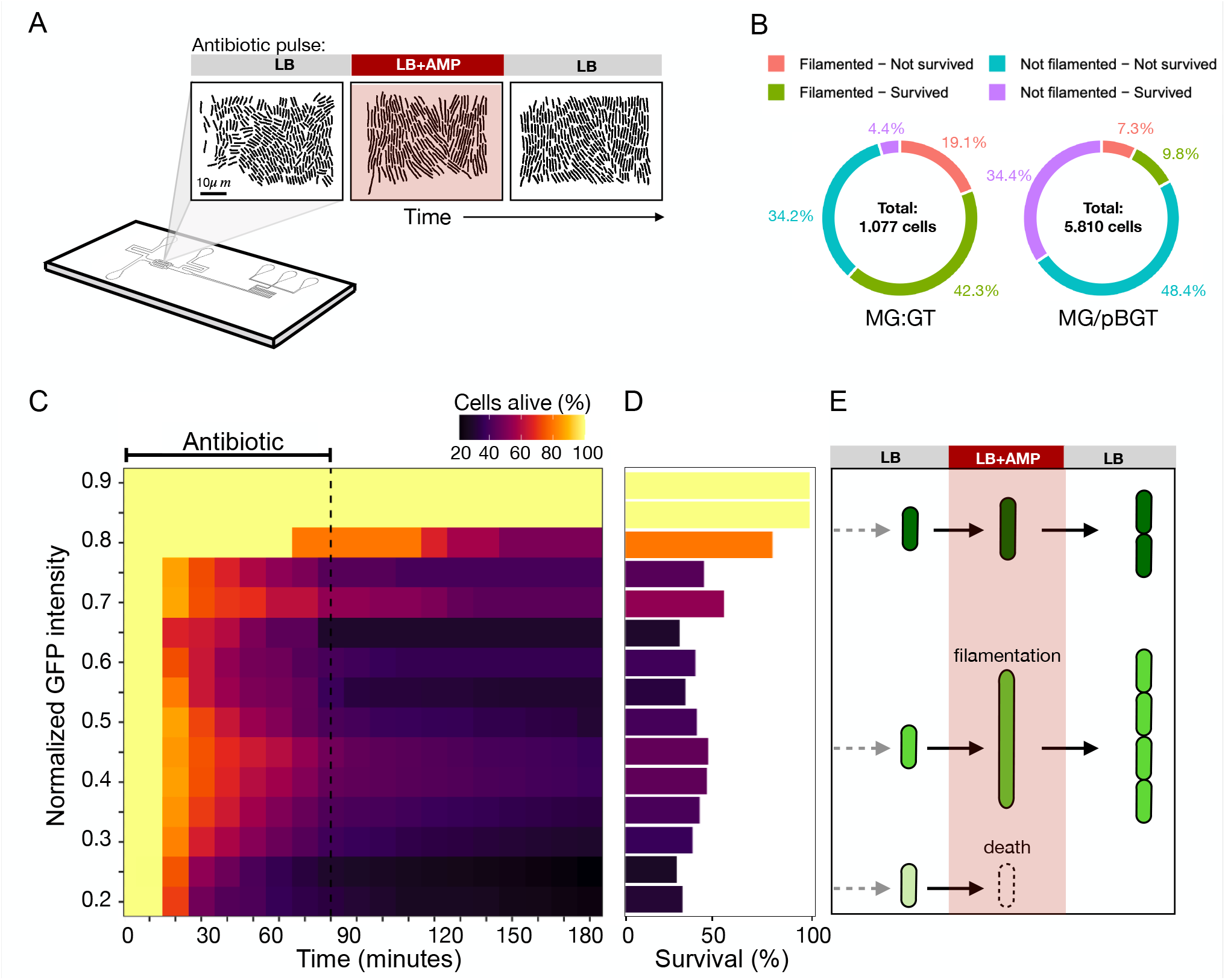
Single-cell analysis of a semi-lethal pulse. A) Schematic diagram illustrating a microfluidic experiment exposing populations of MG:GT and MG/pBGT to a semi-lethal antibiotic pulse. B) Summary of results obtained after tracking individual cell lineages in time-lapse movies. Cell lineages are classified based on whether they were able to survive treatment (in green and purple, for cells that produced filaments or not, respectively) or if they died during drug exposure (in red cells that filamented but died, in light blue cells that were killed without triggering the stress response). C) Fraction of cells alive as a function of time for lineages present when the antibiotic was introduced. Y-axis denotes the initial fluorescence of cells in each initial GFP bin, and each box represents the proportion of cells that are still alive in each time step (high survival rates in a light color) D) Histogram of GFP expression in a population of MG/pBGT cells estimated at the end of the microfluidic experiment. The size of each bar represents the probability of survival estimated for each GFP level after exposure to a semi-lethal pulse of AMP. Note how the distribution appears bimodal, with high survival rates at intermediate and very high fluorescent intensities. E) Diagram illustrating that this bimodal distribution is a consequence of a stress response mechanism that produces filamented cells and provides transient resistance to ampicillin in cells with intermediate fluorescent values. Cells with low GFP values before drug exposure have a low probability of survival, while cells with high fluorescent intensities are highly tolerant to the antibiotic.

AMP-induced cell lysis was estimated by staining the medium with rhodamine and measuring the accumulation of fluorescent dye. After a recovery period in drug-free medium, cells were classified according to whether they died or survived the semi-lethal pulse. As the antibiotic concentration and duration of treatment were determined separately for each strain, we expected the semi-lethal pulse to kill approximately half the population. In line with this prediction, only 46.7% of MG:GT cells and 44.2% of MG/pBGT cells survived the antibiotic pulse (Figure 4B).

A retrospective analysis of surviving and non-surviving cells revealed that surviving cells had an elevated duplication rate, measured as the time elapsed between cell division events (87.09 and 106.7 minutes, respectively; Figure S6; p-value*<* 0.005). Similarly, surviving cells had a higher rate of elongation (changes in cell length between consecutive frames) than cells that were killed (Figure S7; p-value*<* 0.005). These results suggest that an enhanced probability of survival is a consequence not of reduced metabolic activity, but of heterogeneity in *bla*_TEM-1_ expression.

Changes in the fraction of surviving cells as a function of GFP expression before drug exposure are shown in Figure 4C. As expected, cells with very high GFP expression had a high probability of survival (54% survival for the top quartile), whereas the mean survival rate for cells in the bottom quartile was below 34%. Interestingly, survival probability was not a monotonously increasing function of GFP intensity, since high survival rates were also observed in cells with intermediate GFP expression (Figure 4D).

### Plasmid-driven phenotypic noise produces a heterogeneous stress response

To investigate if this bimodality in the survival distribution is a consequence of a heterogeneous stress response triggered by a subpopulation of cells, we exposed a MG:GT population to a semi-lethal pulse of AMP and recapitulated the life history of the surviving cells (Figure 5A-B). Note that shortly after being exposed to the antibiotic, some cells ceased dividing but continued to grow, thus producing filaments (see also Figure 4E). Conditional filamentation can be triggered by multiple molecular mechanisms,^58^ including a general stress response – the SOS regulatory network – that regulates the expression of over 50 genes involved in DNA repair, DNA damage tolerance, and the induction of a DNA damage checkpoint that transiently suppresses cell division.^59^

**Figure 5.**
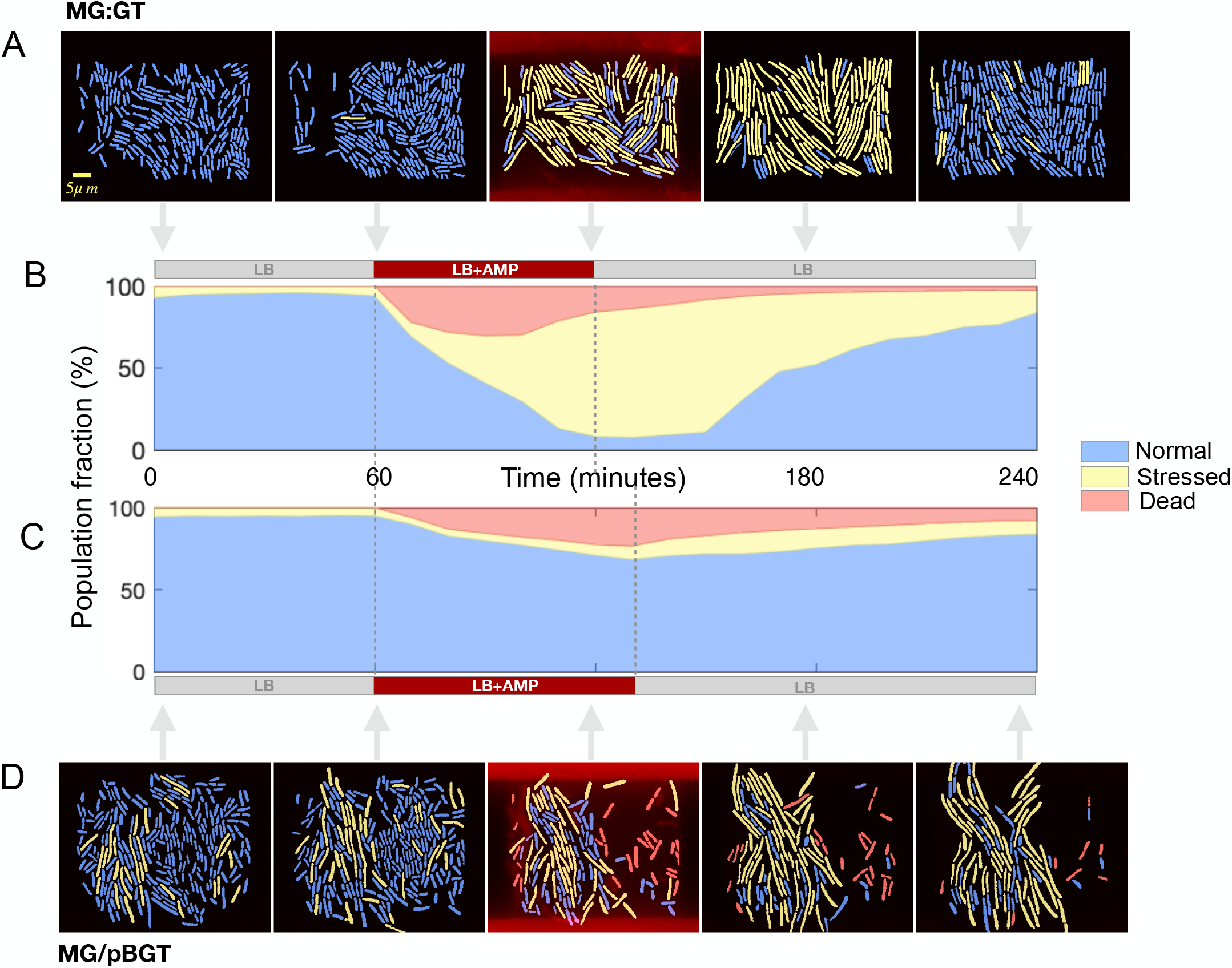
Microscopy montage of a microfluidics semi-lethal pulse. A) Cell classification for a MG:GT population: normal cells (blue), filamented cells (yellow), and dead cells (red). When the antibiotic is introduced into the microfluidic device, MG:GT cells synchronously trigger the SOS response and produce filaments. When the antibiotic is removed, elongated cells resolve and resume normal growth. B) Fraction of the MG:GT population in each cellular state as a function of time (normal cells in blue, stressed in yellow, and dead in red). Most surviving cells exhibit conditional filamentation upon antibiotic exposure and resume normal growth once the drug is withdrawn. C) Fraction of the MG/pBGT population in each cell state. In this case, a smaller fraction of cells produce filaments, as high PCN cells maintain low periplasmic levels of antibiotics and survive without triggering the stress response system. D) Selected frames from a time-lapse movie showing how the plasmid-bearing population responds heterogeneously to antibiotic exposure.

In particular, the SOS response can be triggered by the binding of *β* -lactamase molecules to penicillin-binding protein 3 (PBP3). Lactamase-bound PBP3 acts through DpiBA, a two-component signal transduction system^60^ that induces *sulA*, which in turn inhibits septation by blocking FtsZ polymerization. As a result, cell division is suppressed and bacterial filaments are produced.^61, 62^ Crucially, once the stress is removed, filamented cells reorganize the FtsZ ring, divide, and resume normal growth.^63, 64^

Furthermore, consistent with previous studies,^65^ the temporal expression of genes in the SOS system appeared to be tightly regulated, with 61.4% of cells in the MG:GT population responding synchronously to the antibiotic input and producing filaments (we define a filamented cell as a cell with more than two standard deviations from the mean length of the population before drug exposure). In contrast, the plasmid-bearing population produced a very heterogeneous response, with only 17.1% of cells producing filaments (Figure 5C-D). This was expected, as we have established that variability in PCN maintains a subpopulation of cells that overproduce *β* -lactamase and hence avoid triggering the stress response by maintaining a low periplasmic AMP concentration. Conversely, cells with low PCN are killed by the antibiotic before they can trigger the SOS response.

Histograms of GFP fluorescence in cells of each subpopulation before the introduction of AMP was introduced into the microfluidic device are shown in Figure S8. As expected, the MG:GT population exhibited low variance, with no significant differences in mean GFP intensity detected between subpopulations. In contrast, the plasmid-bearing population exhibited a GFP intensity distribution with high variance. We classified each cell according to whether it was killed or survived drug exposure and according to whether or not the stress response was triggered. Surviving cells in the MG/pBGT population, either had high fluorescence intensity and did not trigger the SOS response (a consequence of increased *β* -lactamase synthesis), or had intermediate GFP fluorescence and survived antibiotic exposure by elongating and delaying cell division.

We also performed an exploratory data analysis, which showed that while PCN (measured by proxy through GFP intensity) is important for cell survival, so is cell length at the moment of the environmental perturbation (see PCA plot in Figure S9). This analysis confirmed that cells with increased survival are small cells with high GFP fluorescence, or cells that were already filamented when exposed to AMP (Figure S10). Our data suggests that plasmid-driven phenotypic noise produces random conditional filamentation, thus enabling the population to adapt to a rapid increase in drug concentration.

### High levels of antibiotic resistance are unstable in the absence of selection

In our microfluidics data, the mean time elapsed between cell duplication events was significantly different between the two strains (36.6 minutes for MG:GT and 88.2 minutes for MG/pBGT; p-value*<* 0.001; Figure S11). Similarly, at the population-level, a comparison of growth rate in strains with different PCNs with respect to plasmid-free cells revealed a negative correlation between growth rate and mean PCN in the absence of selection for plasmid-encoded genes (*R*^2^ =0.562; Figure S12). The cost associated with bearing plasmids is well-documented,^66–68^ particularly for ColE1-like plasmids,^69, 70^ and has been reported for multiple plasmid-host associations in a wide range of bacterial species.^45, 71–73^

The burden associated with plasmid carriage is highly variable and depends on the interaction between plasmids and their bacterial hosts.^50^ This fitness cost can be ameliorated through mutations in genes located either on the chromosome or the plasmid.^74–77^ In addition to these compensatory mutations, another strategy to ameliorate the burden of carrying high-copy plasmids is to reduce the number of plasmids carried per cell. For instance, a previous experimental evolution study reported that mutations near the origin of replication generated a 10-fold amplification in mean PCN, but at a very high fitness cost that resulted in high levels of antibiotic resistance being unstable in the population once the antibiotic was removed.^41^

To assess how rapidly PCN amplification is reversed once the antibiotic is withdrawn, we performed a three-season serial dilution experiment in which a MG/pBGT population was exposed to fluctuating selection (season 1, drug-free; season 2, 32 *mg/ml* AMP; season 3, drug-free). The GFP fluorescence distribution was recorded at the end of each season (Figure 6A). In the presence of AMP, the GFP fluorescence distribution shifted to high expression but rapidly returned to the original fluorescence distribution once the antibiotic was removed. This effect was also observed with high-copy plasmids (Figure S13).

**Figure 6.**
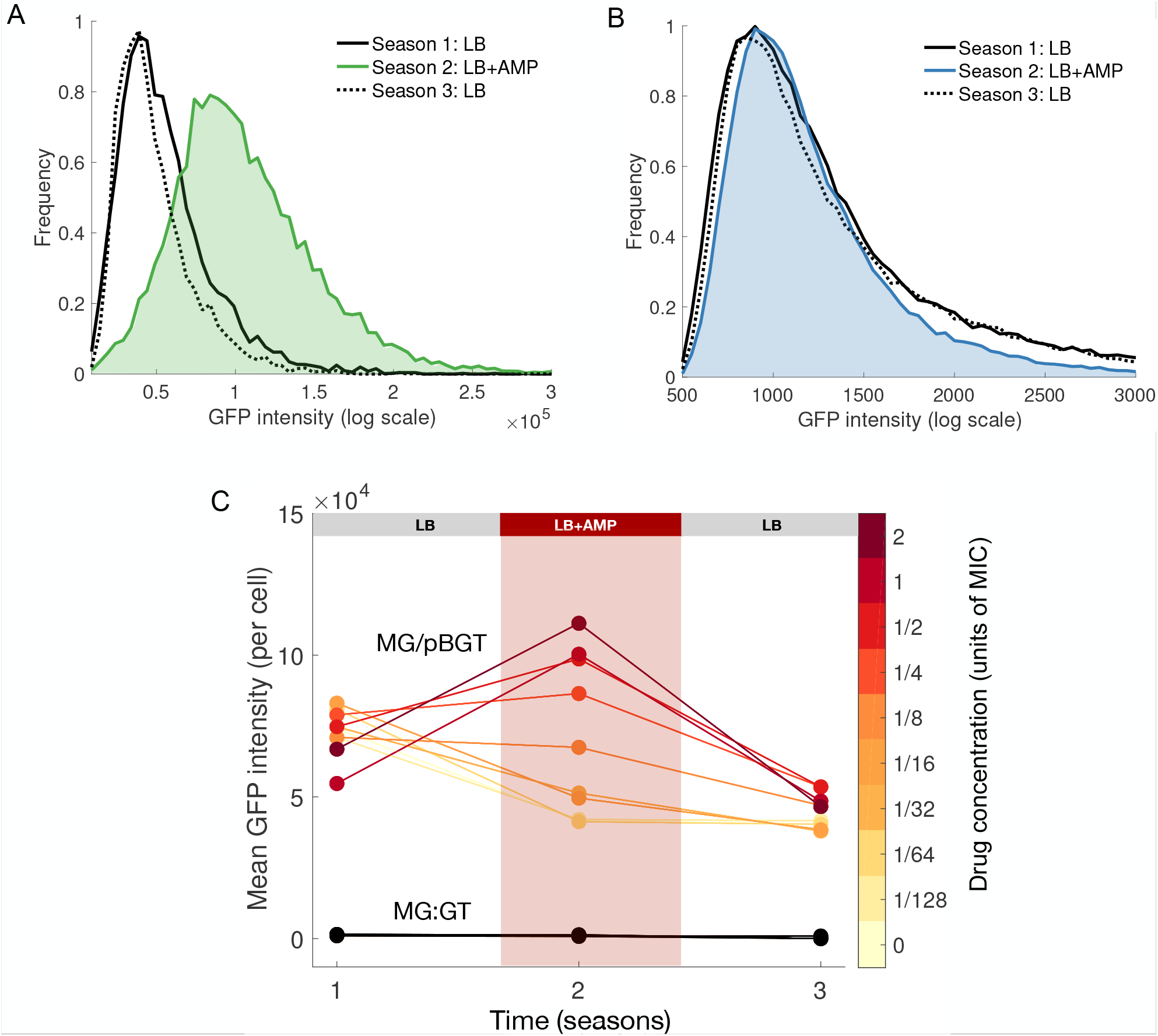
Adaptation to fluctuating environments with different strengths of selection. A) GFP histogram in a population of MG/pBGT exposed to fluctuating selection (Season 1 (LB): solid black line, season 2 (LB+AMP): green area/line, season 3 (LB): dotted black line). Note that the antibiotic shifts the GFP distribution to the right (green area) and is later restored when the antibiotic is removed. B) GFP histogram for MG:GT reveals that GFP distributions coincide independently of the environmental drug concentration. C) Increase in mean fluorescence in the presence of antibiotics is correlated with drug dose (darker red, higher drug concentrations). Once the antibiotic is removed, mean GFP intensity is restored to pre-exposure levels. The black line shows that fluorescent intensity for MG:GT remains constant during the experiment.

Repeat runs of the experiment with different drug concentrations revealed that mean GFP fluorescence of the MG/pBGT population increased proportionally to the strength of selection, and the shift towards higher copy number cells was rapidly reversed after removing the antibiotic (Figure 6C). In contrast, the GFP intensity distribution in MG:GT cultures was the same independently of the presence of antibiotic in the medium (Figure 6B).

### Stochastic plasmid dynamics promotes heteroresistance in a computational model

To further explore the interaction between the stochastic plasmid dynamics and the strength of selection for plasmid-encoded genes, we used a multi-level computational model that incorporates intracellular plasmid dynamics into an ecological framework (Methods). Briefly, the agent-based model explicitly simulates key cellular processes: cell duplication, resource-dependent growth, antimicrobial-induced death, and random plasmid replication and segregation. Propensities of each process are determined from the concentrations of a limiting resource and a bactericidal antibiotic present in a well-mixed environment.

Figure 7A shows numerical realizations of the agent-based model simulating an exponentially-growing population of cells descended from a parental plasmid-bearing cell. We considered the number of plasmids carried by each cell as a time-dependent variable subject to two main sources of noise: (1) imperfect PCN control,^78^ with plasmid replication occurring in discrete events distributed stochastically over time, and (2) plasmid segregation occurring randomly between daughter cells upon division. A consequence of this stochastic plasmid dynamics is that PCN in any individual cell is highly variable over time and, as the culture is no longer synchronous after a few cell duplications, plasmid-bearing populations exhibit high levels of copy number heterogeneity. This heterogeneity results in a PCN distribution with large variance (Figure 7B).

**Figure 7.**
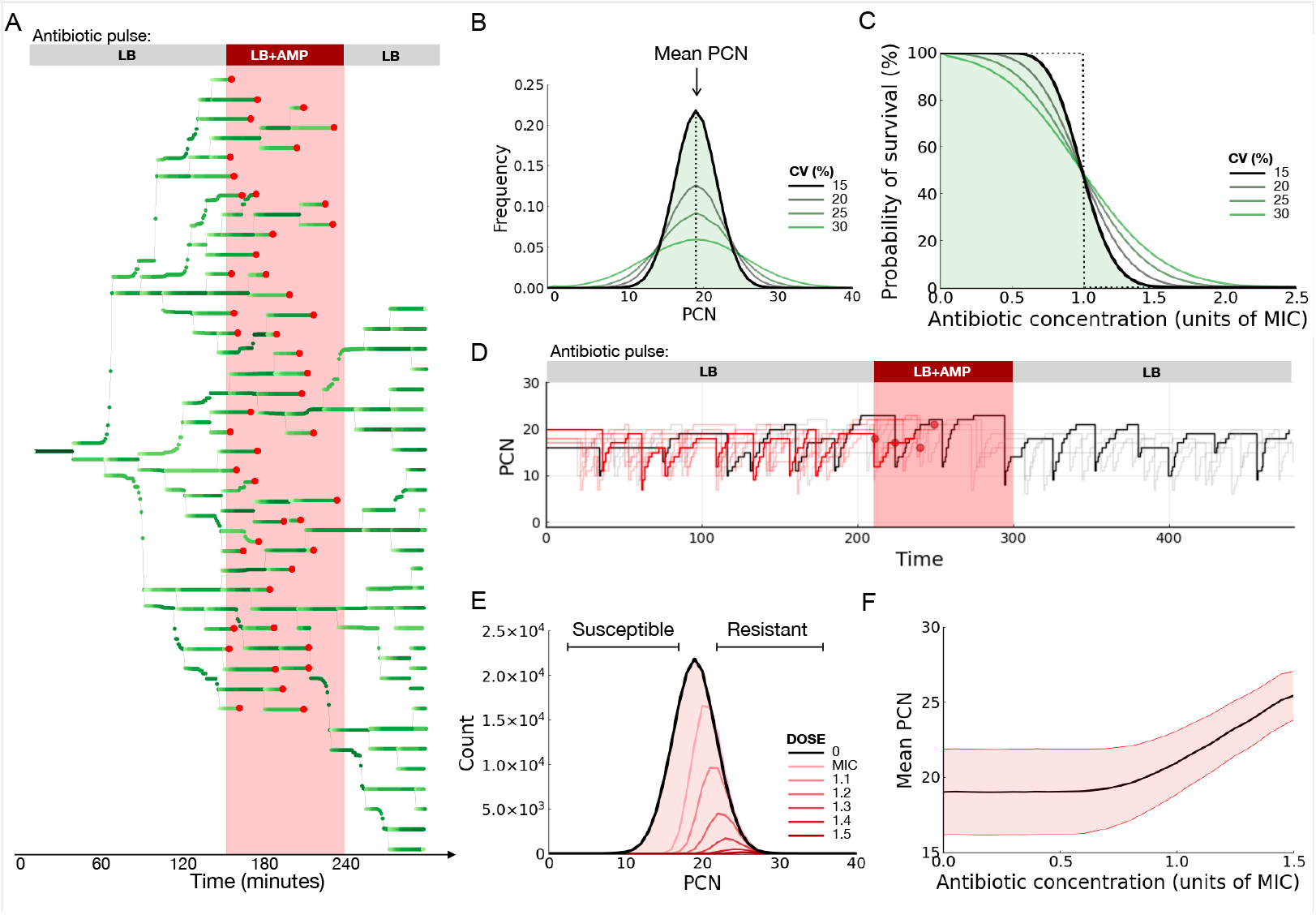
Stochastic plasmid dynamics yield heteroresistant populations. A) Simulations of the plasmid dynamics model of a population growing in drug-free media from a single plasmid-bearing cell. PCN as a function of time is represented in a gradient of greens. The red area represented the time interval when the population was exposed to a lethal drug concentration. Most cells are killed (death events denoted in red), but a few cells carried large PCN during drug exposure and proliferate once the antibiotic is withdrawn. B) Histogram of PCNs estimated using the computational model. The black line denotes the PCN distribution obtained using parameter values described in Table S2, while the green lines illustrate other simulations that produce distributions with larger variance. C) Probability density functions of Normal distributions with a fixed mean and increasing standard deviations. The dotted line illustrates the case when the PCN distribution has zero variance. As the variance of the PCN distribution increases, so does the fraction of cells with an increased probability of survival at high drug concentrations. D) Intracellular plasmid dynamics for individual cells. Lines denote PCN in each cell, with changes in PCN resulting from two random processes: plasmids replicate during cell growth, and segregate between daughter cells during cell division. The red area denotes an interval of drug exposure, and cells killed have trajectories denoted in red. In gray, cells that survived drug treatment and continued segregating and replicating plasmids. E) PCN distributions obtained for different antibiotic concentrations: black line for drug-free environments and, in a gradient of red, the distributions obtained after exposing the heterogeneous population to increasing drug concentrations. F) Mean PCN (black line) and standard deviation (red area) were presented by PCN distributions obtained after exposing the population to a range of antibiotic concentrations. The dotted line illustrates the MIC of the homogeneous population. Note that selection for cells with multiple plasmids occurs even at sub-MIC concentrations.

Based on our experimental data and previous reports,^79, 80^ we assume a linear relationship between PCN and gene dosage. Therefore the probability of an individual cell dying upon exposure to a given antibiotic concentration can be estimated from the number of plasmid copies it carries and the degree of resistance conferred by each plasmid-encoded gene. For instance, if we assume that every cell in the population is equally sensitive to the antibiotic (i.e. a population with low-variance PCN distribution), then we find a drug concentration that kills all cells simultaneously (a dose referred to in the clinical literature as the minimum inhibitory concentration, MIC). Hence the survival probability function of such a homogeneous population is a stepwise function that switches from 1 to 0 at this critical drug concentration (black dotted line in Figure 7C).

However, when we consider a heterogeneous population characterized by a PCN distribution with large variance then, by definition, the population contains cells with fewer or more gene copies than the expected value (green lines in Figure 7B). This implies that the survival probability of heterogeneous populations is lower than that predicted for a homogeneous population at sub-MIC concentrations and higher than the predicted value in high-drug environments (Figure 7C). Indeed, temporal changes in PCN can result in cells with differing degrees of drug susceptibility; as a result, when antibiotics are introduced into the system, only that fraction of cells that had overreplicated the plasmid was able to survive drug exposure (Figure 7D).

In our computational experiments, exposure to antibiotics reduced total bacterial density, but, as cells with low levels of resistance are cleared first from the population, the PCN distribution shifts towards higher values (red lines in Figure 7E). Furthermore, the computational model predicts that the intensity of drug-induced PCN amplification in the population is proportional to the strength of the selective pressure (Figure 7F), with selection for high-copy plasmid cells occurring even at sub-lethal drug concentrations (resulting from killing cells with fewer plasmids than the mean PCN).

Moreover, once the antibiotic was withdrawn, cells that survived continued to grow and divide, therefore randomly replicating and segregating plasmids (see Figure S14). A consequence of this stochastic plasmid dynamics is that cells with low PCN are readily produced through segregational drift. These low-copy cells are at a competitive advantage relative to high-copy sub-populations, and consequently the mean PCN of the population returns to the level observed prior to antibiotic exposure. Similarly with the experimental data, repetition of the computational experiment for different selection strengths revealed that the degree of PCN amplification appears to be correlated not only with the strength of selection, but also with its rate of decay once the antibiotic is removed.

## Discussion

The evolution of antimicrobial resistance in response to the industrialized consumption of antibiotics, specifically those of the *β* -lactam class, is one of the most serious health threats societies face today.^81^ In clinical isolates, heteroresistance can be the result of unstable genomic amplifications,^26^ and has been shown to be the first stage in the progression to *β* -lactam resistance.^82^ Taken together, our data show that cell-to-cell differences in PCN in a clonal population can produce heterogeneity in drug susceptibility in the population, thus enabling plasmid-bearing populations to implement a nonresponsive adaptive strategy that increases their survival in a context of fluctuating selection pressures. Microfluidics uniquely enabled us to connect the plasmid copy number of individual cell lineages (GFP fluorescence) to their phenotypic variability (survival, elongation, or death) under antibiotic pressure and to examine their fate after the antibiotic was removed. Single-cell traces also allowed us to compare our experiments to dynamic computational models.

Moreover, the combination of high-throughput fluorescence measurements with single-cell and population-level susceptibility assays enabled us to show that PCN distribution is modulated by the strength of selection for plasmid-encoded genes, rapidly increasing the mean resistance of the population during selective conditions. Our analysis focused on non-conjugative, multi-copy plasmids that are usually carried at around 10-30 copies per cell; however, plasmid-driven phenotypic noise is not exclusive to high-copy plasmids.^79^ A recent study showed that a conjugative, low PCN populations (1-8 copies per cell) also exhibited large copy number heterogeneity that resulted in noisy expression of plasmid-encoded genes.^83^

We also found that PCN heterogeneity promoted variability in the SOS system, a stress response mechanism that is known to increase resistance to heavy metals^84, 85^ and antimicrobial substances.^86, 87^ This stress response is also known to increase genetic variation^88^ by promoting bacterial mutagenesis^89, 90^ and enabling the horizontal transmission of virulence factors^91^ and antibiotic resistance genes.^92^ The SOS system also produces bacterial filaments, which have been shown to be an adaptive trait with many benefits,^58^ including the promotion of tissue colonization^93^ and increased tolerance to cell wall damage produced by the antibiotics used in this study.^64, 94^

Our study, combined with previous reports, shows that having a phenotypically diverse population is an effective adaptive strategy to survive fluctuating environmental conditions.^95–97^ Transitions between phenotypic states can result from promoter noise;^98^ asymmetry in the cell division process;^99^ or stochastic fluctuations in the concentrations of proteins, mRNAs, and other macromolecules present at low-copy numbers in the cell.^100–102^ We proposed that the stochastic nature of plasmid replication and segregation also produces heterogeneous populations, in which a minority of cells that carried more copies of a plasmid encoding the antibiotic resistance gene *bla*_TEM-1_ survived exposure to a lethal AMP concentration.

Upon removal of the antibiotic from the environment, surviving cells continued growing and dividing, therefore replicating and segregating plasmids. As a result, low PCN cells with increased competitive fitness relative to the highly-tolerant subpopulation emerged, therefore restoring drug susceptibility and compensating for the cost imposed by bearing multiple plasmid copies. Altogether, these results indicate that multicopy plasmids provide a platform for implementing a reversible phenotypic tolerance mechanism that rapidly compensates for the burden of carrying multiple plasmid copies.

Furthermore, transient amplification of selective genes encoded in multicopy plasmids may not be exclusive to *bla*_TEM-1_, as similar effects would be achieved by antimicrobial resistance genes encoding efflux proteins or other drug-modifying enzymes.^8, 103–8, 105^ Other systems where gene dosage is relevant and that scale with gene copy number may also use multi-copy plasmids as platforms for fine-tuning gene activity.^106, 107^ A recent study showed that precise control of gene expression in genetic engineering and synthetic biology can be achieved by tuning PCN in individual cells.^80^ This provides a promising tool for the optimization of synthetic circuits, but also represents a novel approach that can be used for the design of rational treatment strategies that are effective at suppressing heteroresistant populations.

## Acknowledgements

We thank S. Brom, J. Stavans, D. Romero, P. Padilla, M. Ackermann and R. Beardmore for useful discussions and comments on earlier versions of this manuscript. We are also thankful with A. Saralegui from Laboratorio Nacional de Microscopía Avanzada for assistance using the flow cytometer and J. Escudero for the gift of the strains. We also thank the LABNALCIT-UNAM (CONACYT) for technical support using the cell sorter. JCRHB was a doctoral student in Programa de Doctorado en Ciencias Biomédicas, Universidad Nacional Autönoma de México, and received fellowship 59691 from CONACYT. BAL is a student in Programa de Doctorado en Ciencias Bioquímicas, Universidad Nacional Autönoma de México and received fellowship 886346 from CONACYT. JVS received a scholarship from PAPIIT-UNAM (grant IN209419). ASM is funded by the European Union’s Horizon 2020 research and innovation program (ERC grant agreement no.757440-PLASREVOLUTION). JR-B is supported by a Miguel Servet contract from Instituto de Salud Carlos III (ISCIII; grant no. CP20/00154), co-funded by ESB, ‘Investing in your future’. RPM and RCM were supported by a Newton Advanced Fellowship awarded by the Royal Society (NA140196). AFH and RPM were supported by PAPIIT-UNAM (grants IA201418 and IN209419, respectively). This project was also funded by CONACYT Ciencia Básica (grant A1-S-32164) awarded to RPM.

## Materials and Methods

### Bacterial strains and culture conditions

In this study, we used *Escherichia coli* K12 MG1655 bearing a ColE1-like (p15a) plasmid, pBGT, encoding for the *β* -lactamase resistance gene *bla*_TEM-1_ that confers resistance to ampicillin, an *eGFPmut2* gene under an arabinose inducible promoter, and the *araC* repressor. Mean PCN=19.12, s.d.= 1.53.^45^ As a control, a strain *E. coli* K12 MG1655 was used, carrying the *pBADg f p*2, *araC*, and the *bla*_TEM-1_ integrated into the chromosome through the *λ* -phage. Strains bearing plasmid variants G54U and G55U contained a point mutation in the origin of replication: G to U changes at positions 54 and 55 of the RNAI placed in the loop of the central hairpin and affect the RNAI-RNAII kissing complex that controls plasmid replication and PCN. All experiments were conducted in Lysogeny Broth-Lenox (LB) (Sigma-L3022) supplemented with arabinose (0.5% w/v) and appropriate ampicillin concentrations were supplemented as indicated in each experiment. Arabinose stocks solutions were prepared at 20% w/v by diluting 2 g of arabinose (Sigma-A91906) in 10 ml DD water sterilized by 0.22 *µ*m filtration. AMP stock solutions (100 mg/ml) were prepared by diluting ampicillin (Sigma-A0166) directly in 0.5% w/v arabinose LB.

### Antibiotic susceptibility determination

The minimum inhibitory concentration (MIC) of different strains was calculated using dose-response curves performed in 200 *µ*L of liquid media. 96-well plates (Corning CLS3370) supplemented with LB (0.5% w/v arabinose) and a logarithmically-separated range of drug concentrations were used. Antibiotic plates were inoculated from a master plate using a 96-pin microplate replicator (Boekel 140500). Inoculation plates were prepared by adding 200 *µ*L of overnight culture into each well and incubating at 37 ^°^C with 200 rpm shaking. Optical density measurements were performed using a BioTek ELx808 Absorbance Microplate Reader at 630 *nm*. MIC was determined when the reader was unable to detect bacterial growth (2, 32, 43, and 46 *mg/mL* for strains MG:GT, pBGT, G54U, and G55U, respectively).

### Plasmid copy number determination

PCN per chromosome was determined using quantitative polymerase chain reaction (qPCR) with a CFX96 Touch Real-Time PCR Detection System. Specific primers were used for the *E. coli*’s *dxs* monocopy gene as chromosomal reference (dxs-F CGAGAAACTGGCGATCCTTA, dxs-R CTTCAT-CAAGCGGTTTCACA) and primers for the *bla*_TEM-1_ plasmid-encoded gene (Tem-F: ACATTTC-CGTGTCGCCCTT, Tem-R: CACTCGTGCACCCAACTGA) both with amplicon sizes 100 bp as previously described.^45^ In short, samples were prepared following a previously published protocol:^108^ 100 *µl* culture samples were centrifuged at 16,000 g for 60”, the supernatant was removed, and the pellet was resuspended in an equal volume of MilliQ water. Then, samples were boiled at 95 ^°^C for 10’ using a thermoblock and stored at -20 ^°^C for later use. Primers were diluted in TE buffer at 10 *µ*M and stored a -20 ^°^C. Primers’ final concentration was 300 nM. qPCR reactions were performed using SYBR Select Master Mix (Applied Biosystems -4472908) in 96-well flat-bottom polystyrene microplates (Corning 3370) sealed with sterile optical film (Sigma-Aldrich Z369667-100EA). Amplification was performed by an initial 2 min at 50°C activation, then an initial denaturation for 2 min at 95 ^°^C, following 40 cycles of 15 sec denaturation at 95 ^°^C, 1 min annealing, and 1 min extension at 60 ^°^C. After the amplification, a melting curve analysis was performed by cooling the reaction to 60 ^°^C and then heating slowly to 95 ^°^C. PCN was determined using the ΔΔ*C*_*T*_ method.^109^

### Flow cytometry

GFP fluorescence distributions were calculated using imaging flow cytometry in an Amnis ImageStream Mark II by Luminex. INSPIRE software was used to control the machine and acquire data. GFP fluorescence was excited at 488 nm using 75 mv intensity. Data files were processed using IDEAS 6.2 software to only take into account cells on focus using area, aspect ratio, and side scatter features. Files were exported to text files and analyzed with custom scripts in Python and MATLAB. Fluorescence-activated cell sorting of the MG/pBGT strain using a BD FACSAria. An overnight culture was grown on 20 ml of LB 0.5% w/v arabinose at 30 ^°^C, and 200 rpm was sorted into subpopulations. Four subpopulations were categorized by fluorescence intensity and SSC-area features. DNA extraction of sorted subpopulations was made as previously described for qPCR and stored at -20 ^°^C for later use. Plasmid copy number measurements were performed in each subpopulation to evaluate the association between copy number and fluorescence intensity.

### Fitness costs determination

To determine competitive fitness in the absence of antibiotics, each strain was cultured in a 96-well plate with LB supplemented with arabinose 0.5% w/v. A Synergy H1 microplate reader was used to obtain the growth kinetics of each strain by inoculating a 96-well plate with an overnight culture of each strain and growing at 37 ^°^C for 24 hours, reading every 20 minutes, after 30 seconds of shaking. Maximum growth rate estimates were obtained by fitting the mean optical density of N=8 using the R package *GrowthRates* using non-parametric smoothing splines fit.^110^

### Semi-lethal pulse in bacterial populations

Strains of MG::GT and MG/pBGT were exposed to a three-season serial transfer experiment using 96-well plates (8 replicates per strain). An initial inoculation plate was made by putting 200 ml of overnight culture per well. Season 1 (LB) was inoculated from an inoculation plate using a microplate pin replicator. Season 2 (LB-AMP) was inoculated from season 1 after 12 hours of growth. We used the following ampicillin gradient: 0, 1*/*128, 1*/*64, 1*/*32, 1*/*16, 1*/*8, 1*/*4, 1*/*2, 1, and 2 MIC units. In season 3, cultures were transferred to a new LB plate after 12 hours of growth, allowing bacteria to grow for another 12 hours. Plates were sealed using an X-Pierce film (Sigma Z722529) perforating every well to avoid condensation and grown at 37 ^°^C inside a BioTek ELx808 Absorbance Microplate Reader. Measurements were taken every 20 minutes, after 30 seconds of linear shaking at 567 cpm (3 mm). At the end of each season, end-point fluorescence intensity was measured using a BioTek Synergy H1 using OD (630*nm*) and eGFP (479.520*nm*). Plates were then stored at 4^°^C before imaging flow cytometry was performed the following day. A complete independent four-replicate experiment was performed for each strain. DNA samples were extracted at the end of each season to quantify PCN.

### Population-level survival assay

Strains were grown in an LB+Amp media in a 96-well plate under a concentration of AMP determined based on the MIC of each strain. For each LB+AMP plate, we considered 88 populations growing in antibiotics and 8 without antibiotics as controls. Inoculated plates were incubated in a BioTek ELx808 absorbance microplate reader at 30^°^C, with optical density measurements (630*nm*) obtained every 30 min, after 1 min of shaking. After each read, plates were taken out, and a plate sample was taken with a microplate replicator to inoculate a new LB plate. Samples were taken every 30 min, from 0 to 8 hours, then at 18 and 24 hours. New plates were grown in a static incubator at 30 ^°^C for 24 hours. Growth was measured using OD (630nm) and eGFP (479,520 nm) in a Synergy H1 microplate reader after 5 min shaking. An additional experiment was performed for the MG:GT and MG/pBGT strains sampling every 2 hours from 0 to 12 hours and a final sampling at 24 hours.

### *β* **-lactamase inhibitor experiment**

For the *β* -lactamase inhibition assay, sulbactam (Sigma-S9701) was used. First, the ampicillin concentration was fixed to be that of the MIC of MG:GT (2 *mg/ml*). Then, a sulbactam dose-response experiment with MG/pBGT was performed and found that the minimum sulbactam concentration achieved that complete growth suppression was 256 *µ*g/l. Critical AMP and sulbactam concentrations were used to performed a population-level survival assay consisting on exposing 8 replicate populations to fluctuating selection: LB→ LB+AMP+sulbactam →− LB. Samples of four replicates were used for flow cytometry, and the remaining four replicate samples were used for PCN quantification.

### Single-cell microfluidics

A microfluidic device built-in PDMS (polydimethylsiloxane; Sylgard 04019862) from molds manufactured by Micro resist technology GmbH using soft photolithography (SU-8 2000.5) was used for this study. In particular, a micro-chemostat that contains two media inputs and 48 rectangular chambers (40×50×0.95*µm*^3^).^111^ Each confinement chamber traps approximately 1, 000 cells in the same focal plane, enabling us to use time-lapse microscopy to follow thousands of individual cells in time. Chips were fabricated by pouring PDMS into the mold before baking it for 2 hours at 65 ^°^C. Solid chip prints were cut, punched, and bound to a glass coverslip using a plasma cleaner machine (Harrick Plasma -PDC-001) at full power for 1 min and 15 sec. Then we baked them again overnight at 45 ^°^C to ensure binding. Moreover, for each strain, MG/pBGT and MG:GT200, a 1 l titration flask was inoculated with 200*µ*l of an overnight culture (LB at 30 ^°^C and 200 rpm) when the culture reached 0.2-0.3 OD630; it was split into 4 falcon tubes and centrifuged for 5 min at 7, 000 rpm. Supernatant was disposed of, and cells were resuspended by serial transfers into 5 ml of fresh media supplemented with arabinose 0.5% w/v. This dense culture was used to inoculate the microfluidic device. Data acquisitions started 5 hrs after the device chambers were filled and cells were growing exponentially.

After 60 minutes of growth, we switched the environment from LB to LB+AMP. Drug concentration was determined independently for each strain (2*mg/ml* and 8*mg/ml* for MG:GT and MG/pBGT, respectively). Media and antibiotics were introduced into the microfluidic device using a bespoke dynamic pressure control system based on vertical linear actuators (adapted from^112^). The duration of drug exposure was determined based on the time elapsed before the probability of survival of the population exposed to the MIC is below 50% (a semi-lethal pulse; an exposure of 2*mg/ml* for 60 min for MG:GT, and of 8*mg/ml* for 80 min for MG/pBGT). At the end of the period of drug exposure, the population was transferred to a drug-free environment and grown for 120 min for MG:GT, and 100 min for MG/pBGT. Growth media was supplemented with arabinose at 0.5% and Tween20 (Sigma-P2287) at 0.075%, and filtered with .22*µ*m filters. Experiments were conducted at 30 ^°^C, and the ampicillin media was stained by adding 5 *µ*l and 3 *µ*l of a fluorescent dye (rhodamine, Sigma S1402) in 100*ml* of media used to grow MG:GT and MG/pBGT cells, respectively. This red fluorescent dye allowed us to calibrate media inputs inside the microfluidic device and also worked as a dead-cell marker. Rhodamine stock solution was prepared, diluting the powder in ethanol, and stored at 4 ^°^C.

### Fluorescence microscopy

Microscopy was performed in a Nikon Ti-E inverted microscope equipped with Nikon’s Perfect Focus System and a motorized stage. Temperature control is achieved with a Lexan Enclosure Unit with Oko-touch. The microscope was controlled with NIS-Elements 4.20 AR software. Image acquisition was taken with a 100x Plan APO objective without analog gain and with the field and aperture diaphragms as closed as possible to avoid photobleaching. DIC channel captures were made with a 9v DIA-lamp intensity, red channel (excitation from 540 to 580*nm*, emission from 600 to 660*nm* filter), green channel (excitation from 455 to 485*nm*, emission from 500 to 545*nm*). Exposure times were 200*ms*, 200*ms*, and 600*ms* for DIC, green and red channels, respectively. Multi-channel, multi-position images were obtained every 10 minutes in the following order: Red, Green, Lamp-ON, DIC, Lamp-OFF. We added the Lamp-ON optical configuration to allow the bright-light lamp to be fully powered before acquiring the DIC image, while the Lamp-OFF configuration was added to make sure that the lamp was completely off before capturing the next position.

### Image analysis

Microscopy time-lapse images were analyzed using *µ*J, an ImageJ-Python-Napari image analysis pipeline that implements Deep Learning for image segmentation. In short, the pipeline uses ImageJ macros to arrange and manipulate microscopy images. Image segmentation was performed using DeepCell.^113^ Binary masks were corrected manually using bespoke ImageJ macros. Cell tracking was performed using a nearest-neighbor weighted algorithm coded in Python. Cell-tracking was corrected manually using a custom cell viewer coded in Napari.^114^ Lineage reconstruction was performed in Python, obtaining thousands of single-cell time-series of fluorescent intensity and cell length, as well as time-resolved population-level statistics, including the probability of survival to the antibiotic shock and the distribution of fluorescent intensities. Our cell viewer also allows easy lineage data visualization and plotting. Code used to analyze images is available in a public repository: https://github.com/ccg-esb-lab/uJ/

### Computational model

A stochastic individual-based model was developed, where cells are modelled as computational objects. Each cell may have a specific plasmid copy number derived from a Normal distribution *N*(*µ, σ*) where *µ* is the mean copy number of the population and *σ* stands for the copy number variability. Cells grow by incorporating a limiting resource, *R*, following a Michaelis-Menten function; this function the cost entailed by the number of plasmids. The plasmid cost follows a linear relationship with respect to plasmid copy number. Cells divide when they reach an energy threshold. Upon division, plasmids are segregated randomly (with a probability of 0.5) to the daughter cells. They began to replicate plasmids following a probability determined by 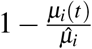, where *µ*_*i*_(*t*) denotes the number of copies of a plasmid at time *t*, and 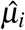 the cell-specific maximum plasmid copy number. The action of the antibiotic is implemented using an individual resistance/susceptibility profile derived from a linear approximation of the experimentally determined population MIC and population copy number, so each cell survival decision was based on their resistance profile, the actual antibiotic concentration, and a random noise modifying this threshold. Numerical experiments of the model were implemented in Julia, with code available in a public repository: https://github.com/ccg-esb-lab/pBGT/

## Supplementary material

Movie S1: MG/pBGT exposed to an antibiotic ramp.

Movie S2: MG/pBGT exposed to a semi-lethal pulse of AMP.

Movie S3: MG:GT exposed to a semi-lethal pulse of AMP.

**Figure S1.**
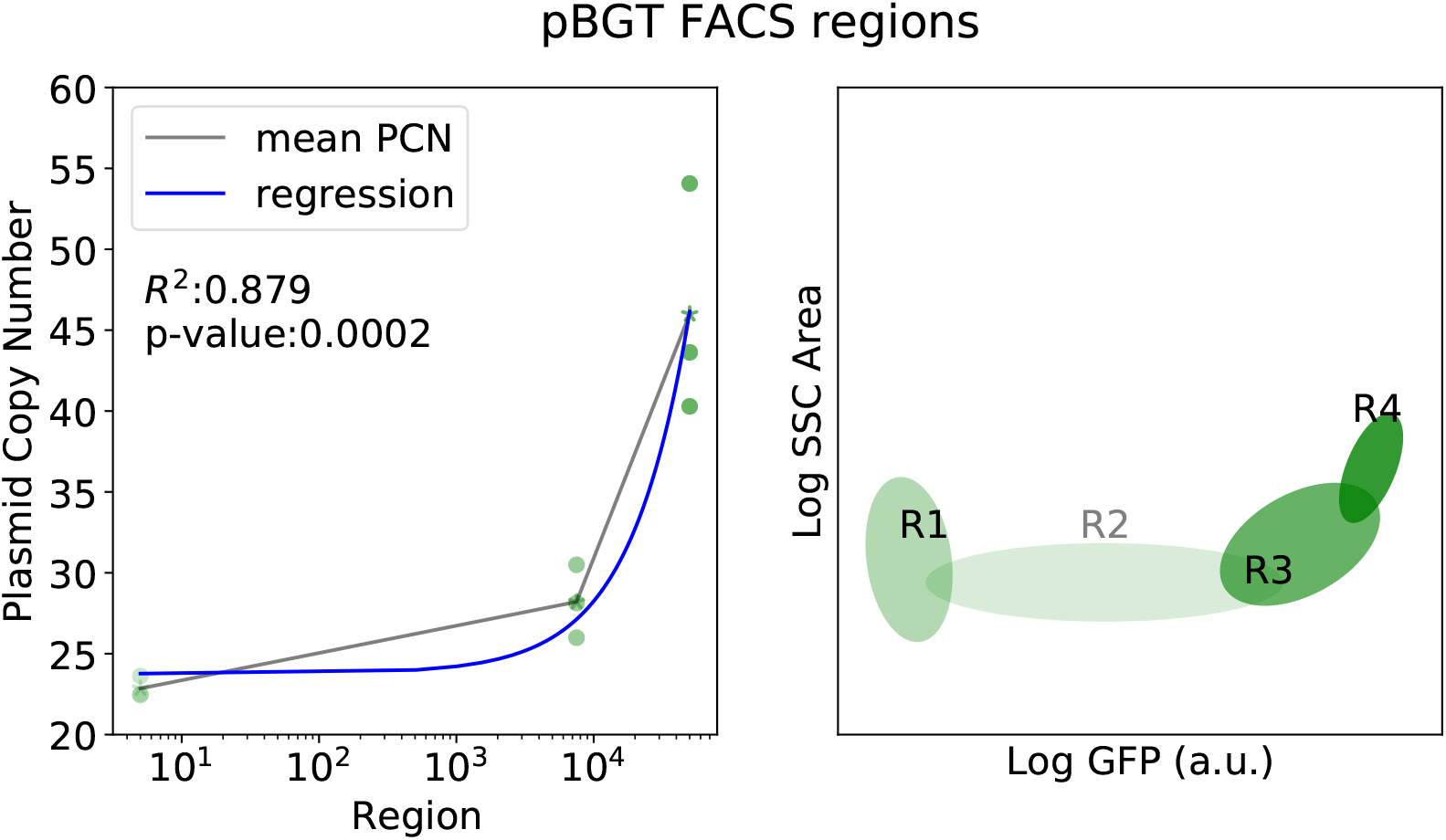
Correlation between PCN and fluorescent intensity. A) Positive correlation between GFP intensity and plasmid copy number estimated using qPCR, for different subpopulations obtained by sorting cells based on their fluorescent intensity. B) Regions used to separate cells based on their fluorescent intensity using a flow cytometer cell sorter.

**Figure S2.**
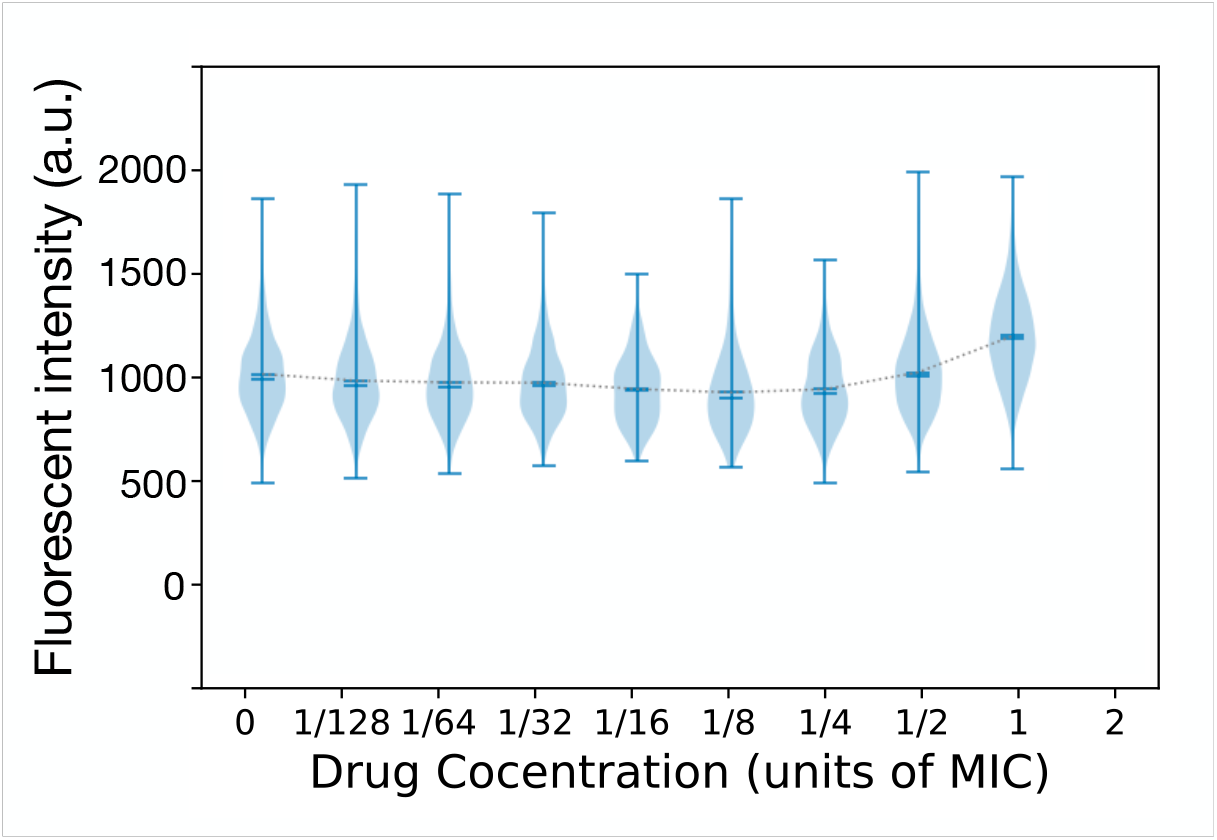
Mean fluorescence of GFP distributions remains constant in populations of MG:GT exposed to increasing concentrations of antibiotic.

**Figure S3.**
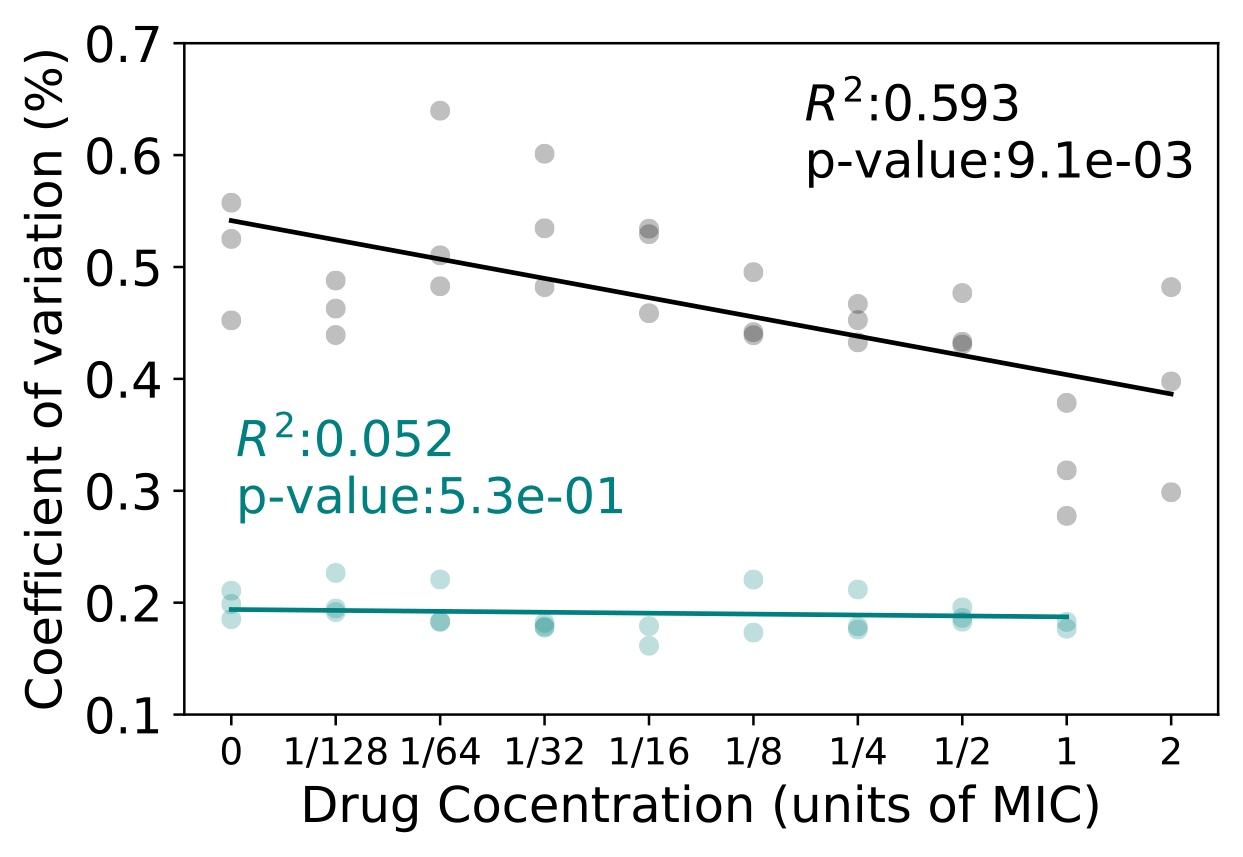
Coefficient of variation of GFP distributions in response to selection. Data for MG/pBGT is denoted with grey circles and for MG:GT in blue circles. Best fit linear regression is shown as solid lines. The Pearson correlation coefficient for the coefficient of variation in pBGT is *R*^2^ = 0.593, suggesting that selection is acting upon the plasmid copy number distribution. In contrast, the coefficient of variation MG:GT remains constant as a function of drug concentration (*R*^2^ = 0.052).

**Figure S4.**
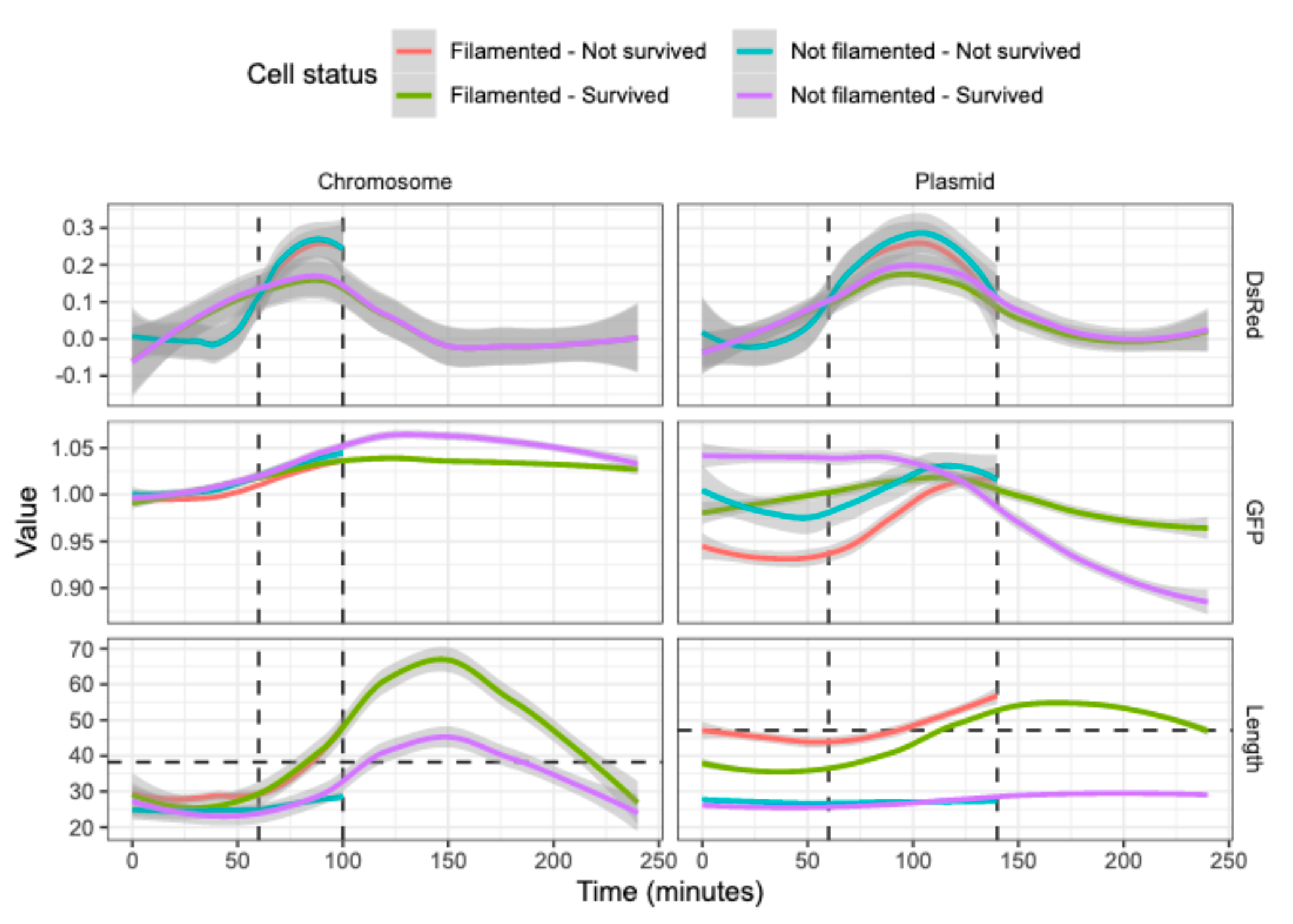
Population-level analysis of microfluidics data. Colored lines represent the average value of each metric as a function of time, with gray shaded area representing the 95% confidence interval. Dotted vertical lines represent the start and end of antibiotic exposure. A) Note a faster increase of DsRed intensity for the non-surviving populations in both experiments, consistent with an increase in the concentration of red fluorescent dye inside the cell. B) For the GFP fluorescent intensity, the chromosomal strain exhibits a stable expression over time, while the plasmid-bearing strain shows a decline in GFP observed for the population that did not survive. C) GFP intensity as a function of time. Note that MG:GT cells that did not filament continued to grow past the filamentation threshold (horizontal dotted line) even after the antibiotic is withdrawn. Eventually mean length of the population reduces as filamented cells resolve and continue growing normally. For the plasmid-bearing strain, note that filamented cells that were killed exhibited a larger cell length than surviving cells when the antibiotic was introduced into the environment.

**Figure S5.**
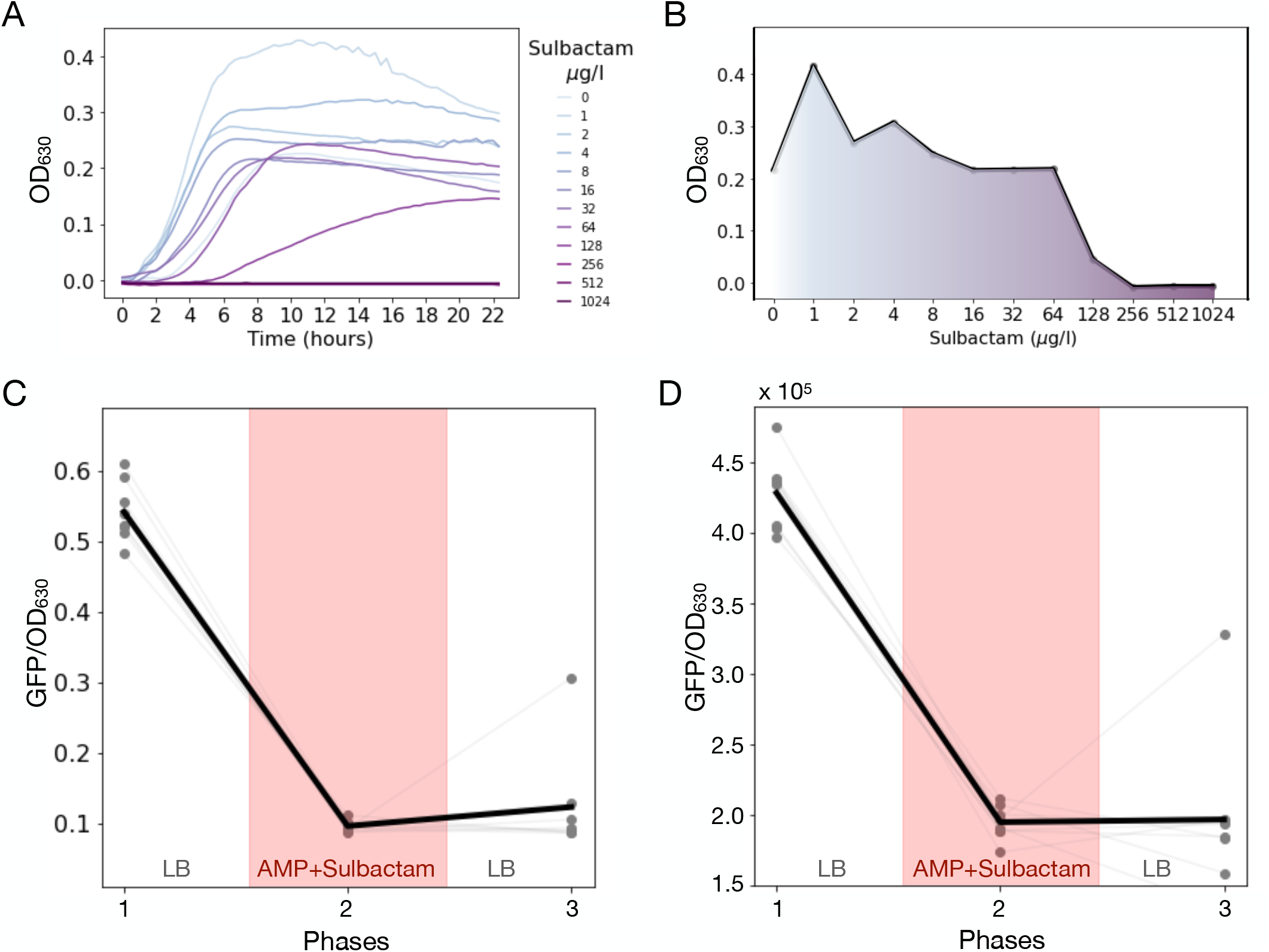
Survival assay with AMP and a *β* -lactamase inhibitor. A) Growth curves obtained for MG/pBGT populations exposed to 2 *mg/ml* of AMP and a range of sulbactam concentrations (low sulbactam doses in light blue, high concentrations in purple). B) Final optical density as a function of sulbactam concentration. We consider that bacterial growth is completely suppressed at concentrations higher than 256*µg/l* of sulbactam. C) Optical density (OD_600_) measured after 12 hours of growth in a 3-season survival assay (season 1: LB; season 2: LB + sulbactam (256*µg/l*) + AMP (2 *mg/ml*); season 3: LB). Gray lines represent different replicates (*N* = 8), with the mean OD_600_ represented with a black line. Of note, only one replicate exhibited growth after the recovery period. D) Normalized fluorescence intensity of populations exposed to a 3-season serial dilution experiment. Note that supplementing the media with sulbactam reduced the relative fluorescent intensity exhibited by the population during the drug exposure period, in contrast to previous experiments performed in the absence of sulbactam, where we observed an increase in fluorescence during AMP exposure.

**Figure S6.**
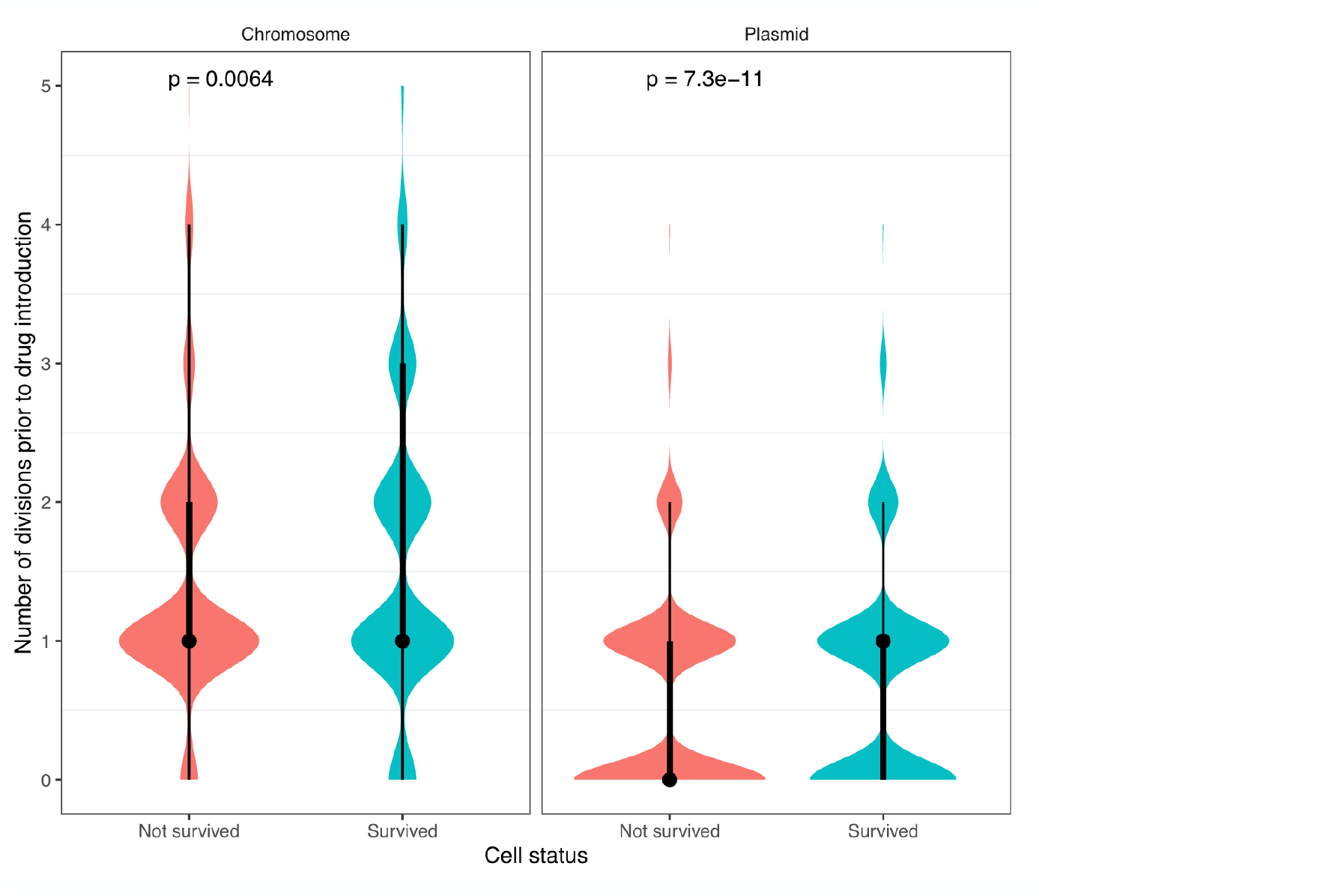
Single-cell duplication rates. Division events were recorded for each cell lineage during the hour prior to drug exposure for MG:GT (left) and MG/pBGT (right). Cells that survived the semi-lethal pulse are denoted in green, and cells killed by the antibiotic in red.

**Figure S7.**
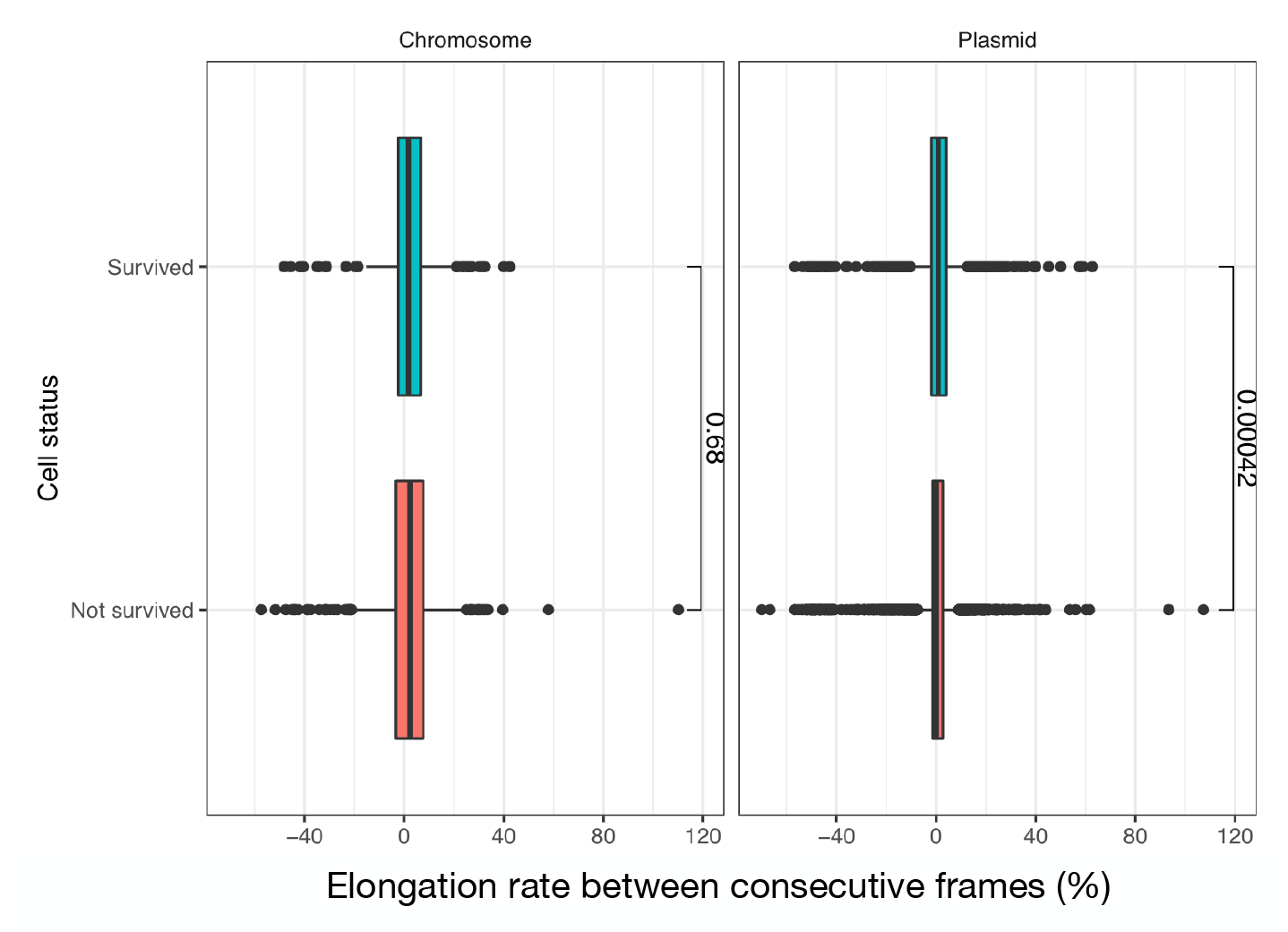
Single-cell elongation rates. Difference in cell length for individual cells in consecutive frames. (left: MG:GT, right: MG/pBGT). In green, cells that survived the semi-lethal pulse and, in red, cells that were killed by the antibiotic.

**Figure S8.**
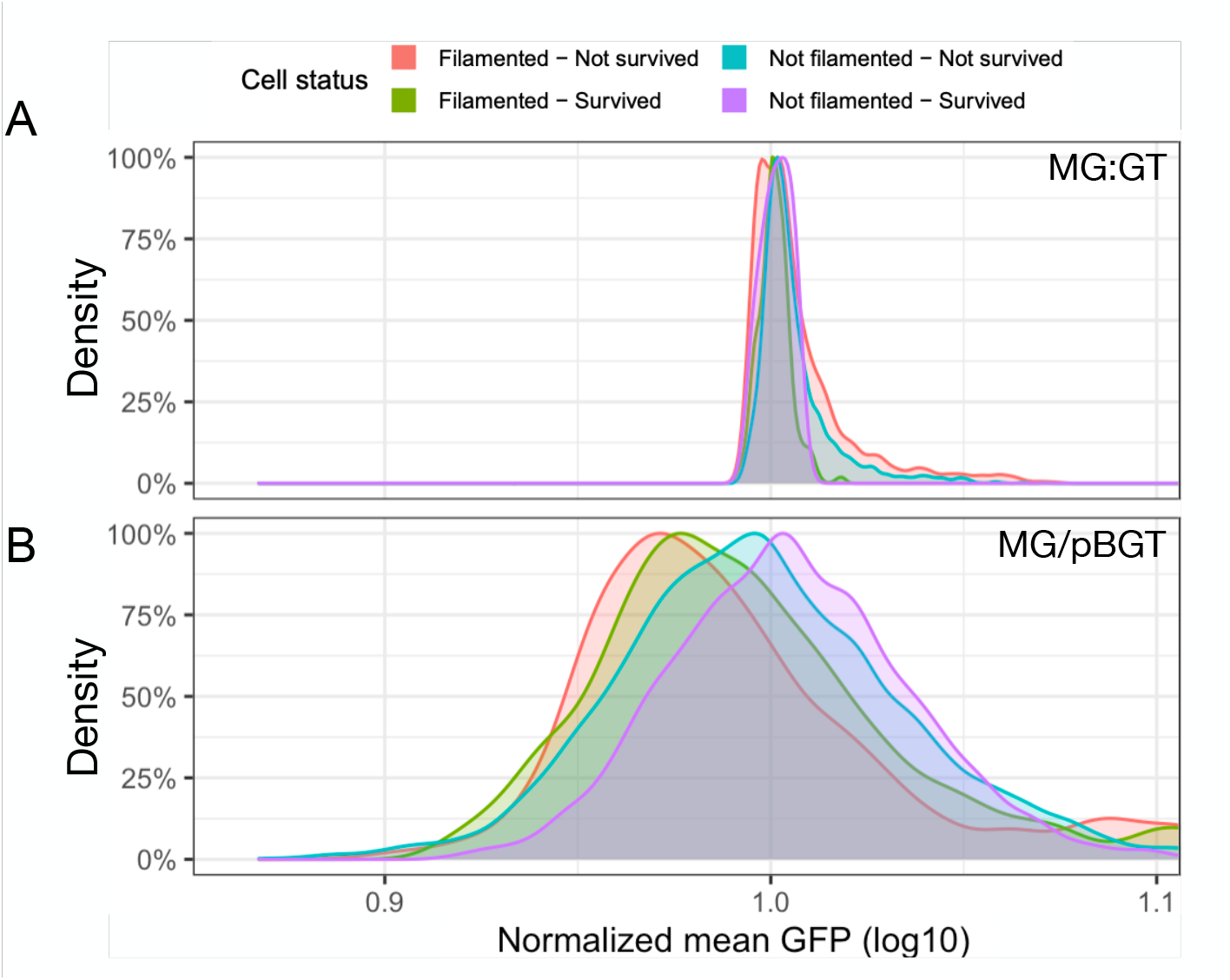
Histograms of fluorescent intensity for classified cells. A) Cells in MG:GT exhibit a fluorescent distribution with low variance and with no significant differences in mean GFP between cells that produced filaments and were killed (red) or survived (green), as well as for cells that did not produce filaments and died (blue), and those that survived drug exposure (purple). B) GFP distributions of the plasmid-bearing population exhibit large variance. Cells that survived showed increased mean fluorescence relative to cells that were killed. For surviving cells, mean GFP was significantly lower for cells that did not produce filaments with respect to cells that triggered the SOS response system.

**Figure S9.**
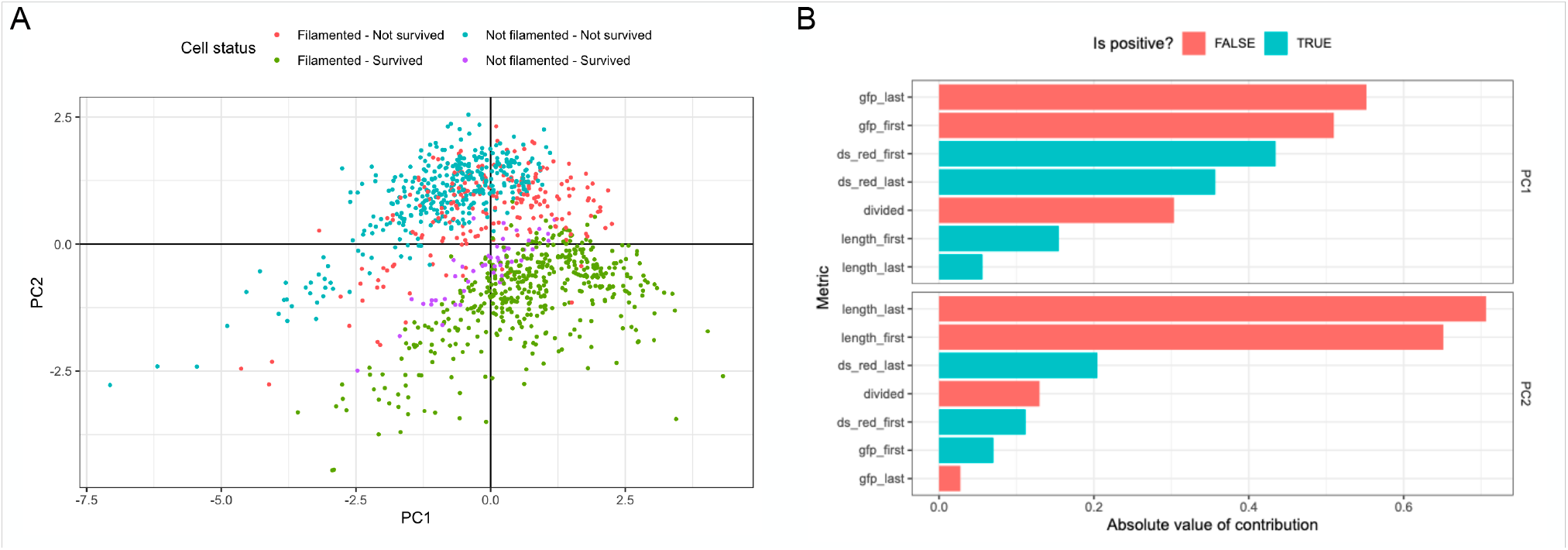
Principal Component Analysis emphasizes the importance of cell length and GFP intensity in cell survival. A) When integrating quantitative information obtained by analyzing time-lapse movies of a semi-lethal pulse, a dimensionality reduction analysis showed a clear separation between the surviving cells and those that were killed by the action of the antibiotic. B) Individual contribution of each variable for the first two components of the PCA analysis. For the first component, the initial and final GFP measurements explained most of the variability. The second component was determined by the length of the cell.

**Figure S10.**
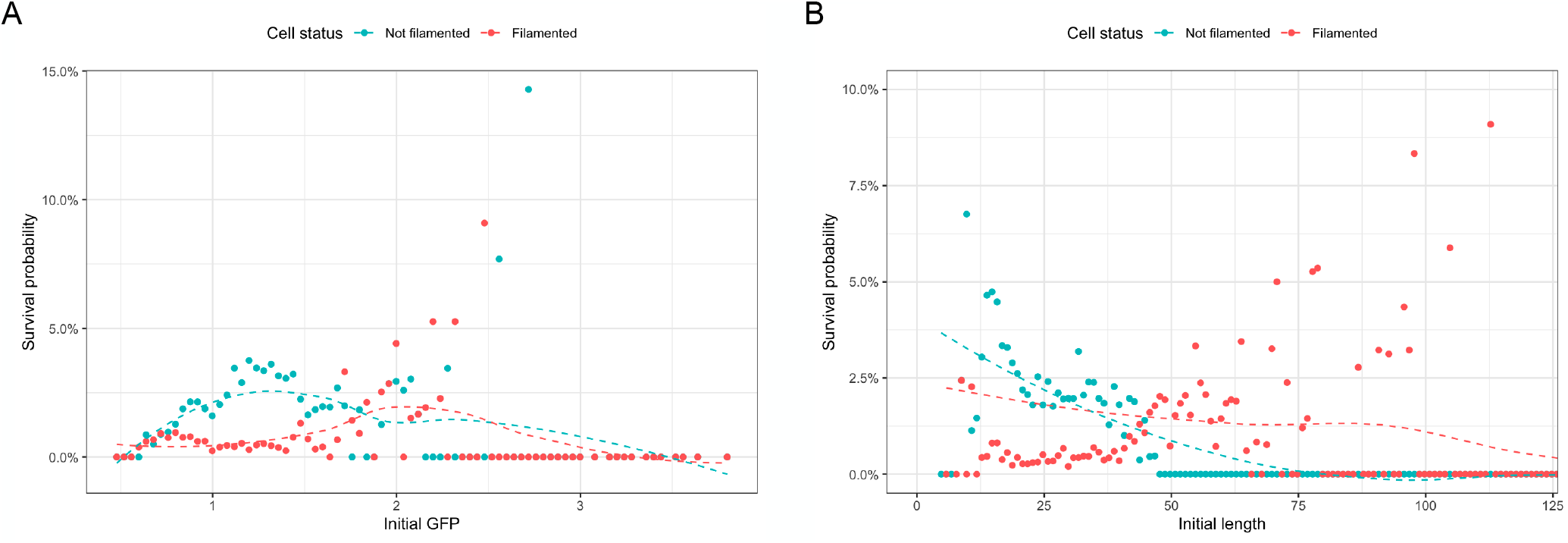
Survival probability of cells with different cell lengths and fluorescent intensities prior to drug exposure. A) Survival of cells that did not produce filaments (blue dots) is maximized at low values of fluorescent intensity. At intermediate GFP levels, a large fraction of surviving cells produced were cells that produced filaments (red dots). B) Cell length at the moment the antibiotic was introduced into the system is an important factor in determining if cells produced filaments or not. At low values of cell length, non-filamented cells exhibited a larger probability of survival than cells that filamented. In contrast, cells that produced filaments exhibited a survival probability that correlates with cell length. Survival probability for very long cells is very low.

**Figure S11.**
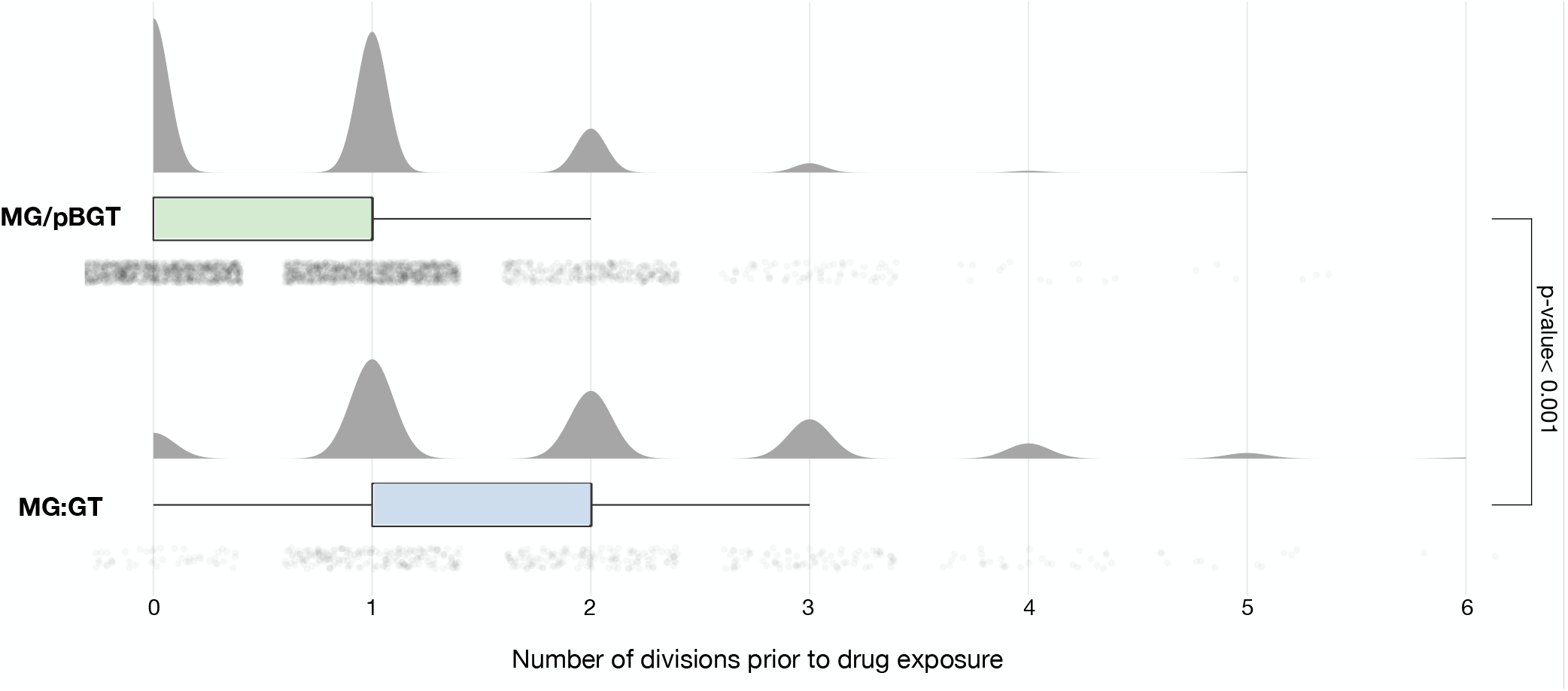
Fitness cost measured in single-cell data. Number of cell divisions before drug exposure for MG/pBGT (green) and MG:GT (blue). Note that the plasmid-bearing strain presented significantly fewer divisions compared to the chromosomal strain, consistent with prior studies showing that carrying plasmids is associated with a fitness cost in non-selective conditions.

**Figure S12.**
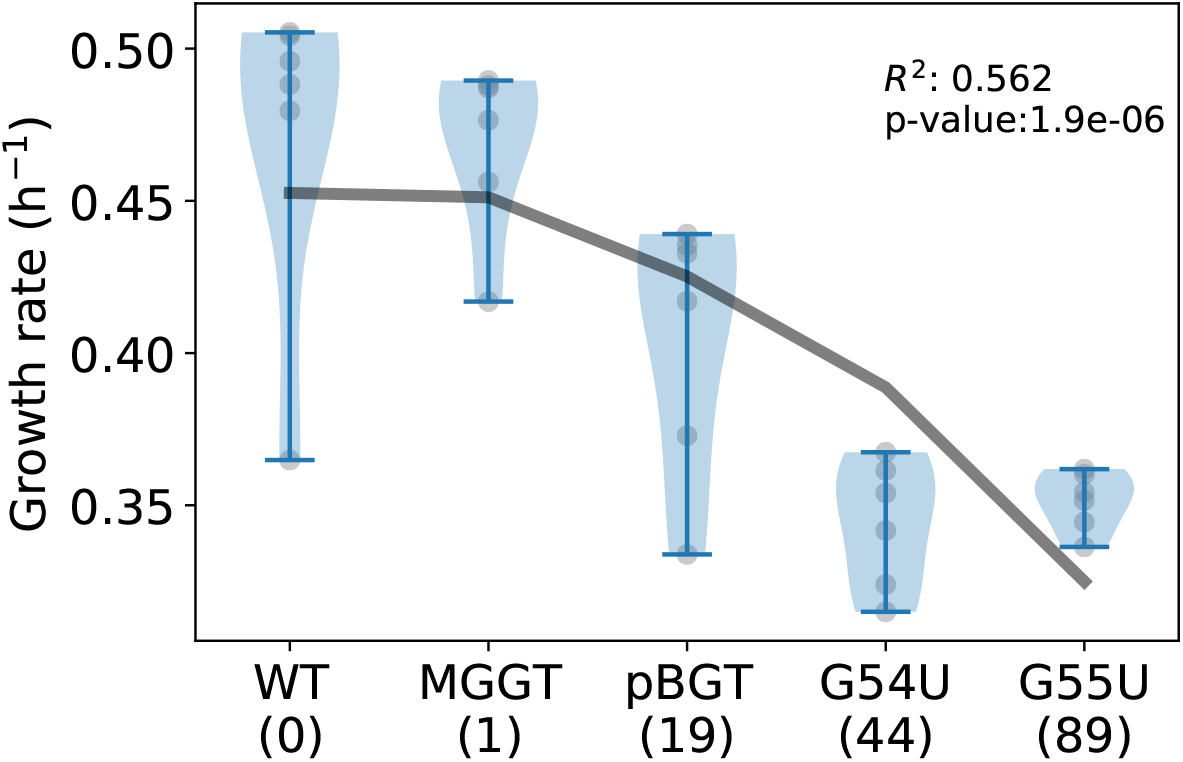
Fitness cost estimated at a population-level. Growth rates for different strains obtained by fitting a growth curve using non-parametric smoothing splines. As expected, there is a negative correlation between fitness in drug-free environments and the number of plasmids carried by each cell. Growth rates ANOVA p-value is 2.91e-07 indicating significant differences. A follow-up Tukey’s Honest Significant Differences analysis yields the following strain pairs with p-value *<* 0.05 : WT-pBGT, pBGT-MGGT, pBGT-G54U.

**Figure S13.**
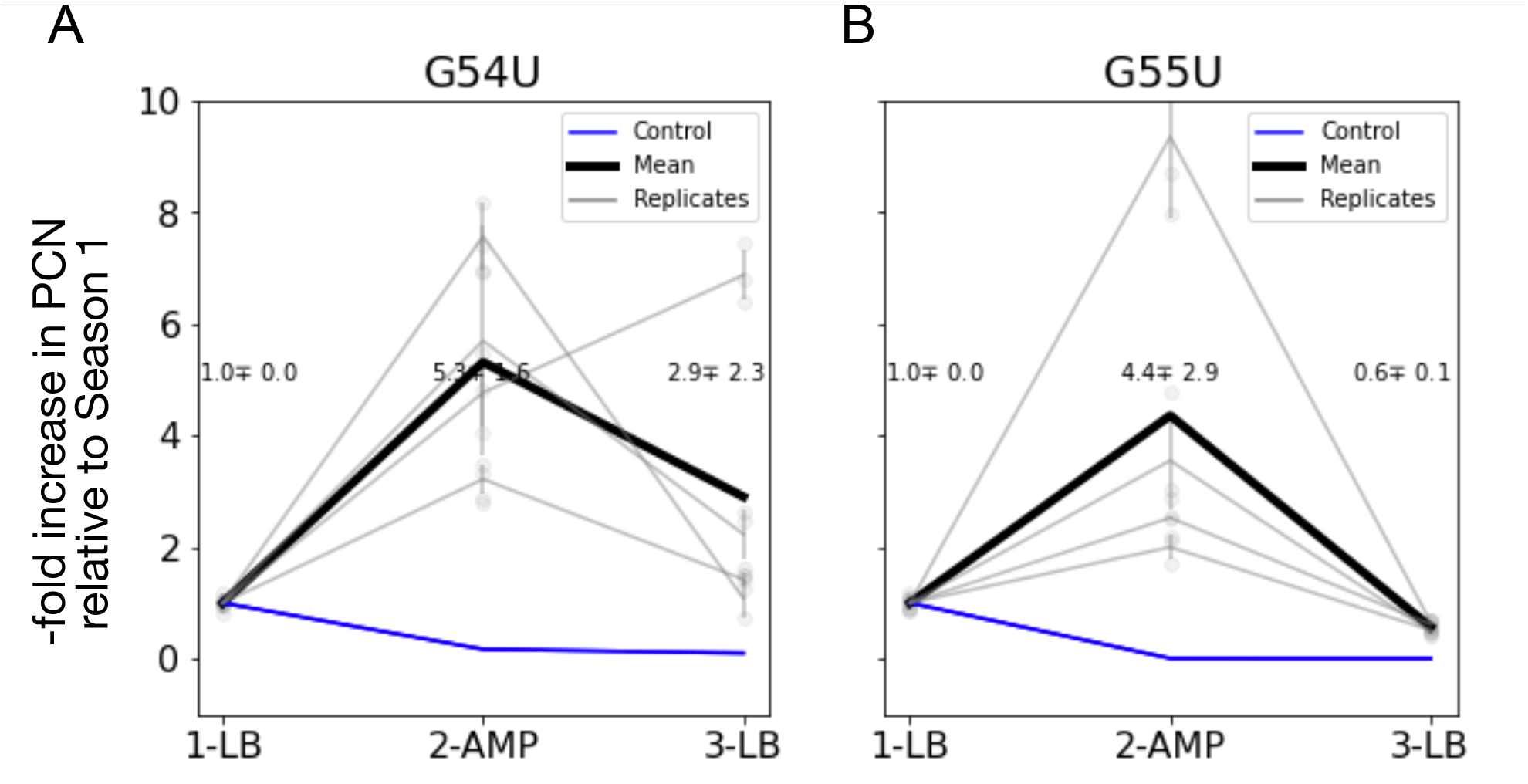
Rapid gene amplification is unstable in high-copy plasmids. A) Fold increase in PCN (relative to season 1) for strain MG/G54U in a three-season serial dilution experiment (black line represents the mean over *N* = 4 replicates, in grey). During the second season, a sub-lethal concentration of AMP is deployed, selecting for high-copy plasmid cells, therefore increasing five-fold the mean PCN in the population. In the third season, the antibiotic is removed and the mean GFP fluorescence intensity decrease to the levels exhibited prior to drug exposure. B) Mean GFP intensity for MG/G55U also shows a rapid increase in fluorescence during drug exposure and a rapid decline once the drug is removed.

**Figure S14.**
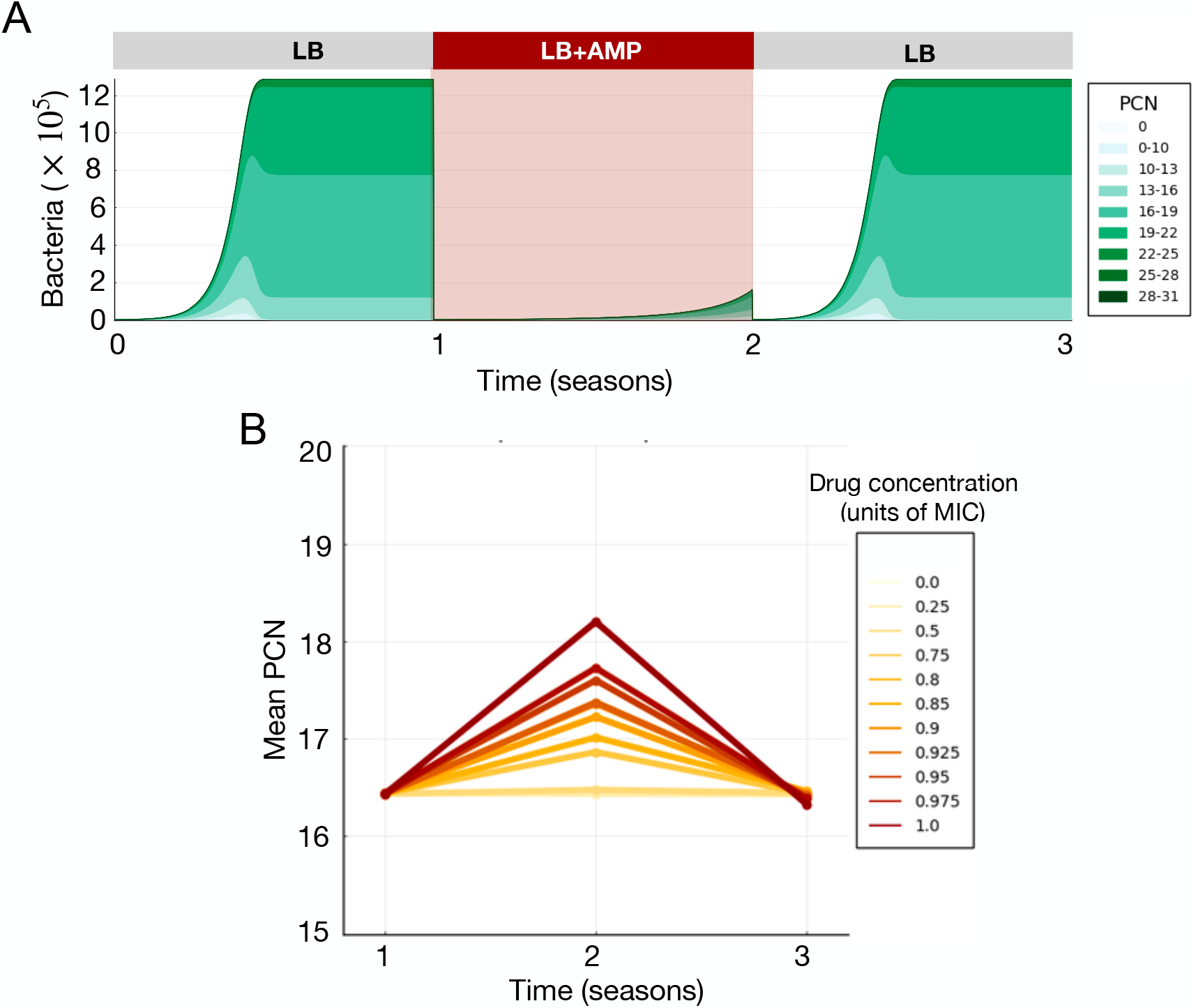
Stability of PCN amplification in the computational model. A) Number of bacteria as a function of time in a three-season serial dilution experiment (with antibiotic deployed in season 2). Stacked areas represent the fraction of cells in the population with different PCNs (plasmid-free in white and increasing PCNs in a range of green). B) Mean PCN at the end of each season in experiments performed with increasing concentrations of antibiotics (low doses in yellow and a lethal dose in red). Note how the increase in mean PCN observed during the selective phase of the experiment is proportional to the drug concentration. The antibiotic was removed in season 3, and the mean PCN exhibited by the population is restored to levels displayed before drug exposure.

**Table S1.**
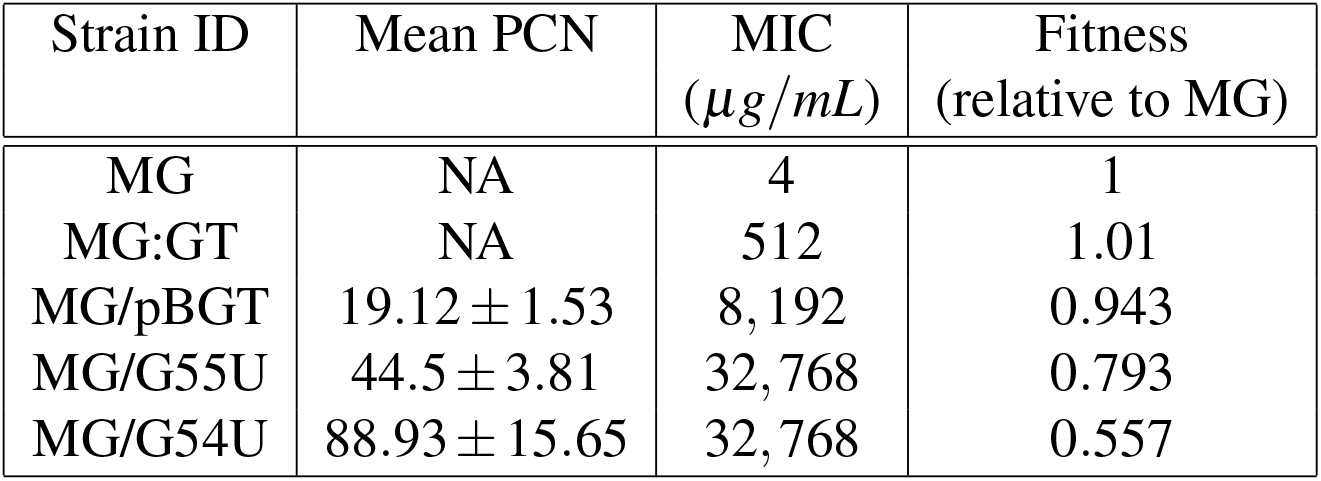
List of *Escherichia coli* MG1655 strains used in this study.

**Table S2.**
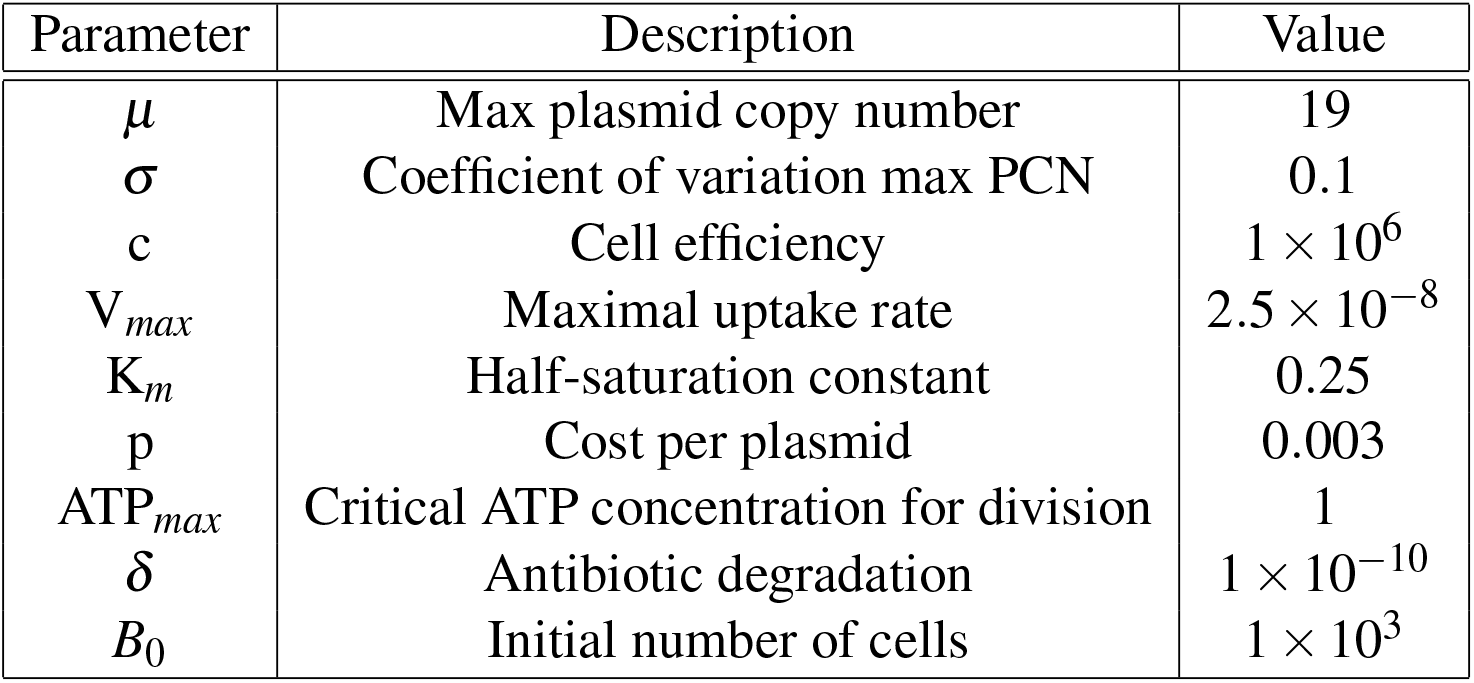
Parameter values used in the numerical simulations of the agent-based model.

| Parameter | Description | Value |
| --- | --- | --- |
| $\mu$ | Max plasmid copy number | 19 |
| $\sigma$ | Coefficient of variation max PCN | 0.1 |
| c | Cell efficiency | $1 \times 10^6$ |
| $V_{max}$ | Maximal uptake rate | $2.5 \times 10^{-8}$ |
| $K_m$ | Half-saturation constant | 0.25 |
| p | Cost per plasmid | 0.003 |
| $ATP_{max}$ | Critical ATP concentration for division | 1 |
| $\delta$ | Antibiotic degradation | $1 \times 10^{-10}$ |
| $B_0$ | Initial number of cells | $1 \times 10^3$ |

## Notes

### Competing Interest Statement

The authors have declared no competing interest.

